# Single-cell transcriptome analysis of human, marmoset and mouse embryos reveals common and divergent features of preimplantation development

**DOI:** 10.1101/385815

**Authors:** Thorsten Boroviak, Giuliano G Stirparo, Sabine Dietmann, Irene Hernando-Herraez, Hisham Mohammed, Wolf Reik, Austin Smith, Erika Sasaki, Jennifer Nichols, Paul Bertone

## Abstract

The mouse embryo is the canonical model for mammalian preimplantation development. Recent advances in single-cell profiling allow detailed analysis of embryogenesis in other eutherian species, including human, to distinguish conserved from divergent regulatory programs and signalling pathways in the rodent paradigm. Here, we identify and compare transcriptional features of human, marmoset and mouse embryos by single-cell RNA-seq. Zygotic genome activation correlates with the presence of Polycomb Repressive Complexes in all three species, while ribosome biogenesis emerges as a predominant attribute in primate embryos, supporting prolonged translation of maternally deposited RNAs. We find that transposable element expression signatures are species-, stage- and lineage-specific. The pluripotency network in the primate epiblast lacks certain regulators operative in mouse, but encompasses WNT components and genes associated with trophoblast specification. Sequential activation of GATA6, SOX17 and GATA4 markers of primitive endoderm identity is conserved in primates. Unexpectedly, OTX2 is also associated with primitive endoderm specification in human and nonhuman primate blastocysts. Our cross-species analysis demarcates both conserved and primate-specific features of preimplantation development and underscores the molecular adaptability of early mammalian embryogenesis.

## Introduction

Metazoan life relies on the ability to develop specialised cell types from a single cell. In mammals, preimplantation development entails timely embryonic genome activation, establishment of a pluripotent cell population to form the foetus and segregation of extraembryonic tissues for successful implantation. This is a highly adaptive process, subject to distinct selective pressures (Sheng, 2015; Friedli and Trono, 2015; Tseng et al., 2017). While the mouse model has been instrumental for our understanding of mammalian development, comparatively little is known about early human and non-human primate embryogenesis.

Primate development is protracted compared to rodents. Human zygotic genome activation (ZGA) occurs around the 8-cell stage, in contrast to the 2-cell stage in mouse (Braude et al., 1988; Yan et al., 2013; Blakeley et al., 2015). At the compacted morula stage, in both rodents and primates, outer cells establish apical-basal polarity providing the basis for segregation of inner cell mass (ICM) and trophectoderm (TE). In human, POU5F1 exhibits prolonged expression in TE (Niakan and Eggan, 2013), in contrast to earlier restriction of Pou5f1 in the mouse ICM. Lineage specifiers NANOG and GATA6 are initially co-expressed in the rodent and primate ICM, and resolve into mutually exclusive patterns (Roode et al., 2012; Niakan and Eggan, 2013; Blakeley et al., 2015; Boroviak et al., 2015; Nakamura et al., 2016; Stirparo et al., 2018), concordant with the segregation of epiblast (EPI) and primitive endoderm (PrE) lineages in the late blastocyst. The EPI represents the founding population of the embryo proper (Gardner and Rossant, 1979), while PrE gives rise to the yolk sac (Artus and Hadjantonakis, 2012; Schrode et al., 2013). In both rodents and primates, embryo implantation into the uterine wall is a landmark event, upon which the EPI acquires epithelial polarity (Enders et al.,1986; Bedzhov and Zernicka-Goetz, 2014) and the regulatory network governing pluripotency is reconfigured (Boroviak et al., 2015; Nakamura et al., 2016).

Rodent and primate embryos implant prior to gastrulation, unlike many other eutherian species, including rabbit, pig, sheep, cow and dog. Upon implantation, the mouse EPI forms an egg cylinder with a pro-amniotic cavity. The amnion is specified via the amniochorionic fold during gastrulation (Arnold and Robertson, 2009; Rossant and Tam, 2009; Pereira et al., 2011). In contrast, primates segregate extraembryonic amnion directly from the EPI after implantation, giving rise to a flat embryonic disc (Rock and Hertig, 1948; Enders et al., 1986; Enders and King, 1988; Enders and Lopata, 1999). Primordial germ cells may be specified from nascent amnion in primates (Sasaki et al., 2016), underscoring the importance of this lineage decision. The ability of primate EPI to form an extraembryonic tissue immediately after implantation is a distinctive feature of primate development, but the underlying transcriptional circuitry at the late blastocyst stage has remained elusive.

Single-cell profiling of human embryos (Yan et al., 2013; Blakeley et al., 2015; Petropoulos et al., 2016) has revealed a multitude of ICM-associated transcription factors, epigenetic regulators and signalling pathway components. However, inherent limitations in the provenance of supernumerary human embryos by the in vitro fertilisation (IVF) route can yield research samples of varying cellular integrity, viability in culture and developmental stage. Despite these challenges, comparison to the mouse ICM has unveiled important differences, including specific expression of KLF17 and ARGFX and increased TGFβ signalling pathway components. However, comparative transcriptional analysis of the second lineage decision and mature EPI specification has been impeded by lack of single-cell RNA-seq data for late mouse ICM samples to resolve distinct EPI and PrE populations (Blakeley et al., 2015). Ultimately, mouse-to-human comparisons alone are unable to elucidate subtle regulatory adaptations between individual species from broader evolutionary features.

Here we constructed a framework for cross-species analysis of embryonic lineages over a time course of preimplantation development in mouse, human and a non-human primate, the common marmoset (*Callithrix jacchus*). We hypothesised that defining hallmarks of early primate development would be consistently observed in human and marmoset, but not in mouse. Compiling stage-matched single-cell transcriptomes from three mammalian species allowed us comprehensive insight into maternal programs, genome activation, stage-specific transcriptional regulatory networks, signalling pathways and transposable element signatures.

## Results

### Cross-species transcriptome analysis of mammalian preimplantation development

Single-cell embryo data was assembled from published studies and newly generated samples, to span a uniform time course of preimplantation development in human, marmoset and mouse (Fig. 1A). For human we used a compendium of RNA-seq data from multiple embryo profiling studies (Yan et al., 2013; Blakeley et al., 2015; Petropoulos et al., 2016) and extracted stage- and lineage-specific transcriptomes (Stirparo et al., 2018) that consistently recapitulate known marker expression *in situ* (Blakeley et al., 2015; Niakan and Eggan, 2013; Deglincerti et al., 2016).

**Fig. 1:**
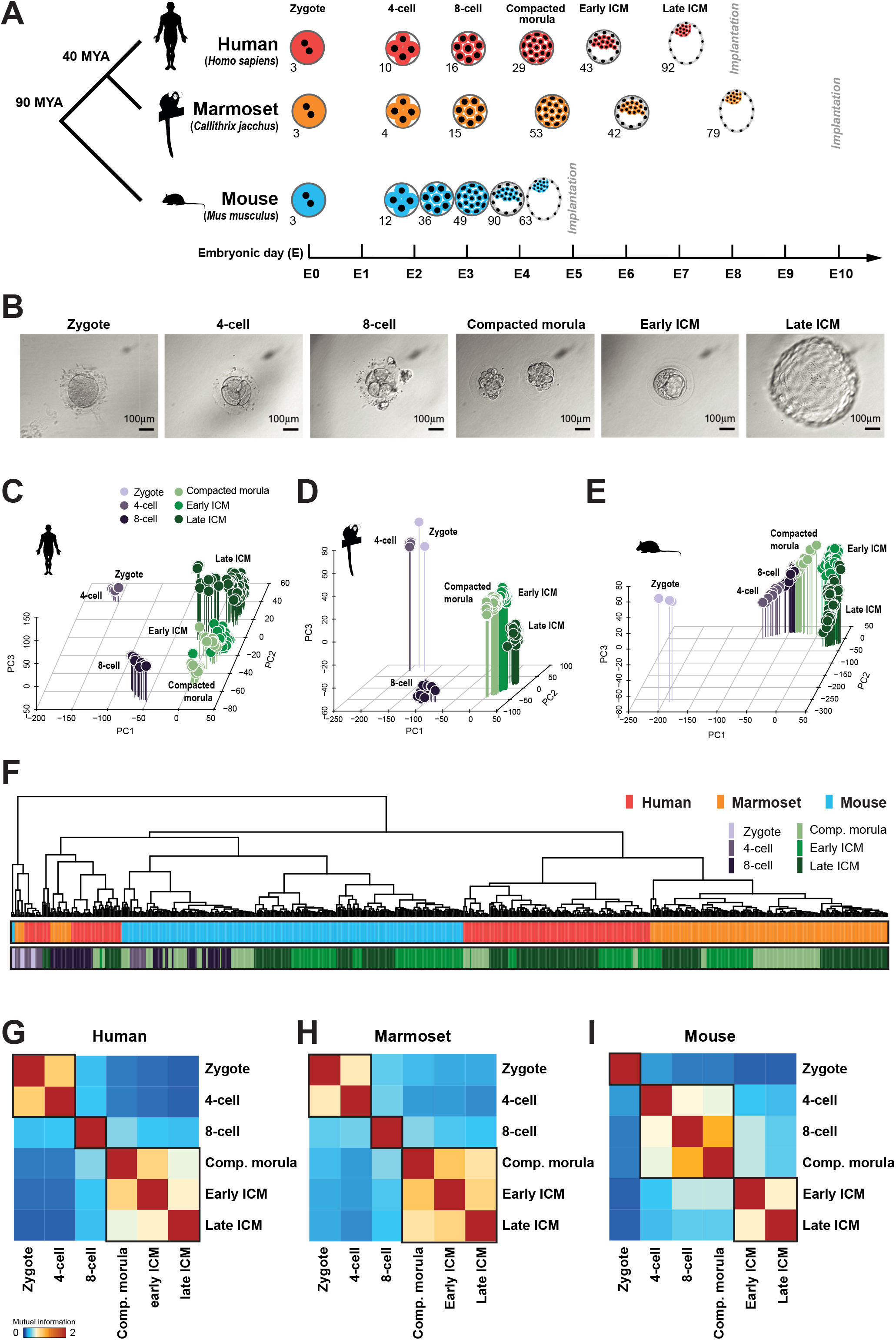
Global analysis of human, marmoset and mouse preimplantation stages. (A) Summary of single-cell RNA-seq data considered in this study. Individual transcriptome numbers are indicated for each developmental stage. MYA = million years. (B) Phase contrast images of marmoset embryos processed for transcriptional profiling. (C–E) PCA of single-cell embryo data for each species (FPKM >0). (F) Pearson correlation distance of preimplantation stages of human (red), marmoset (orange) and mouse (blue), with stages indicated below as in (C). (G–I) Mutual information entropy between preimplantation stages.

We then produced single-cell RNA-seq data from common marmoset embryos developed *in utero*, spanning zygote to late blastocyst preimplantation stages. Non-human primate species provide the means to collect embryos directly by non-surgical uterine flush, improving consistency of sample quality and embryo staging relative to IVF (see Methods). After screening for quality control we retained 196 transcriptomes from 16 embryos over six developmental stages, including early and late ICM cells obtained by immunosurgery (Boroviak et al., 2015) (Fig. 1B, Table S1).

We compiled a similar data series for mouse by combining published data with newly sequenced embryos. To date, the most comprehensive transcriptome profiling analysis of mouse preimplantation development (Deng et al., 2014) did not extend to segregation of EPI and hypoblast (PrE) at the late blastocyst stage, which had limited previous cross-species comparisons to the early ICM (Blakeley et al., 2015). We therefore augmented this dataset with 117 single-cell samples from the early and late ICM (Mohammed et al., 2017).

The complete dataset comprised 642 individual transcriptomes from six preimplantation stages in three mammalian species. Samples were stage-matched to permit direct cross-species comparison, and sequencing libraries for all constituent samples were produced with the Smart-seq protocol (Picelli et al., 2013) to minimise technical differences.

Principal component analysis (PCA) of human (Fig. 1C, S1A), marmoset (Fig. 1D) and mouse (Fig. 1E, S1B) showed tight clustering of samples by developmental stage and time. In mouse, zygotes differed most from other developmental stages (Fig. 1E). However, human and marmoset 4-cell embryos clustered closely with zygotes, while the 8-cell stage was distinct. Significantly, the major ZGA event is reported to occur at the 8-cell stage in human (Braude et al., 1988; Yan et al., 2013). Later preimplantation stages (compacted morula, early and late ICM) from primate were in close proximity to, but spatially segregated from 8-cell embryos (Fig. 1C–E). Separation between zygote/4-cell to 8-cell clusters, and 8-cell to the remaining stages, accounted for most of the variability. This pattern suggests two major transcriptional waves during primate ZGA and contrasts with that observed in mouse, where the greatest separation occurs between zygote and 4-cell, consistent with ZGA at the 2-cell stage (S1B).

Hierarchical clustering of all samples based on orthologous genes showed that the transcriptional state of zygotic embryos interrogated prior to ZGA were most related, regardless of species and distinct from other developmental stages (Fig. 1F, S1C). The remaining mouse samples clustered predominantly with early primate embryos (4-cell and 8-cell), while human and marmoset compacted morulae, early and late ICM were consistently associated. To further assess individual stages we performed correlation analysis based on mutual information entropy (Fig. 1G–I). In human, zygote and 4-cell embryos correlated tightly, and were distinct from the 8-cell stage. Consistent with PCA results, compacted morulae, early and late ICM were co-localised (Fig. 1G). The marmoset closely recapitulated the human profile (Fig. 1H) and late primate stages showed a high degree of similarity in a combined analysis (Fig. S1D). In contrast, developmental stages in mouse followed a different pattern: 4-cell, 8-cell and compacted morulae formed one cluster, while early and late ICM were distinct (Fig. 1I). Thus the global transcriptional program of the preimplantation embryo differs significantly between mouse and primates.

### The primate maternal program is enriched for ribosomal genes and contains a distinct set of epigenetic regulators

Maternally deposited transcripts are abundant in the ovum and persist to varying extents throughout early embryonic development. Among these, maternal effect genes have been defined in mouse as functionally required for early embryogenesis (Kim and Lee, 2014), but such data are not available for primates. Consistent with the established model, 3905 maternal transcripts were robustly detected (average FPKM >=10) in the mouse zygote, with 120 present at high levels (average FPKM >300, Table S2). The majority of these high-abundance RNAs were also found in both primate species, with the notable exceptions of *POU5F1*, *HSF1* and *DICER* (Fig. 2A, B).

**Fig. 2:**
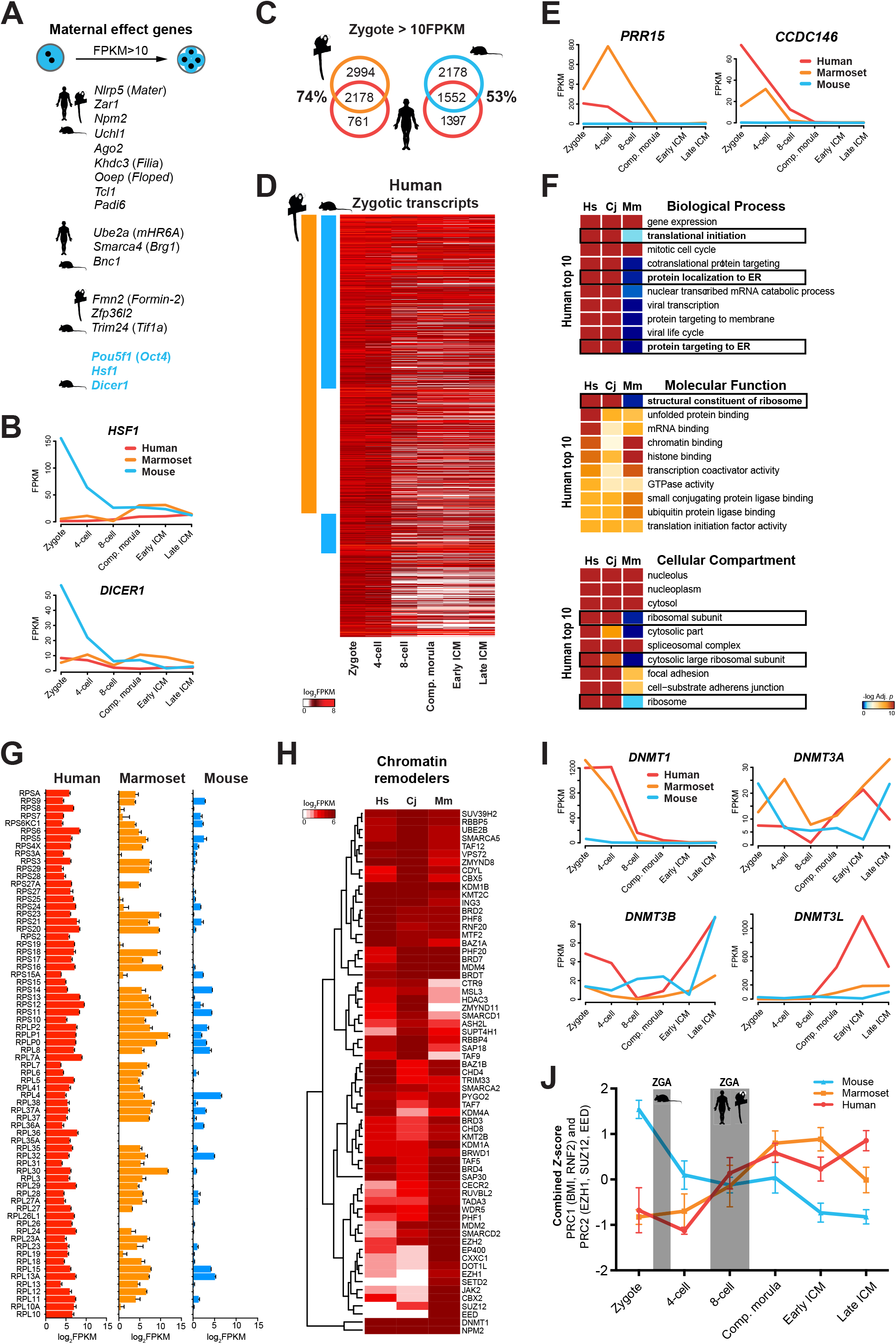
Cross-species analysis of maternal gene transcripts. (A) Schematic of mouse maternal effect genes according to (Kim and Lee, 2014). Symbols indicate transcripts present in the relevant species (FPKM >10). (B) Mouse-specific maternal genes in FPKM. (C) Intersection of maternal transcripts in human, marmoset and mouse zygotes (FPKM >10). (D) Maternal human transcripts (FPKM >10), conserved in marmoset (orange) and mouse (blue). (E) Primate-specific maternal genes in FPKM. (F) GO and pathway significance (-log_10_ *p*-value) ranked according to the top 10 processes statistically enriched in human. (G) Ribosomal transcripts in FPKM. (H) One-way hierarchical clustering of chromatin remodellers in at least one species (FPKM >20). (I) DNA methyltransferases in FPKM. (J) Combined *Z*-score of PRC1 and PRC2 components over developmental time.

To extract the most prominent differences between rodent and primate maternal programs, we compared maternal transcripts in human, marmoset and mouse. Of 2939 detected in human, 74% were conserved in marmoset (Fig. 2C). 53% of human maternal genes were found in mouse, and most displayed decreased abundance over time (Fig. 2D). Conserved maternal factors present in all three species comprised *DPPA3*, *ZAR1*, *PADI6* and *ZAR1L* (Fig. S2A, Table S2). Mouse-specific factors included *Atg5* and the KRAB domain protein-encoding gene *Zfp57*, implicated in imprint protection (Hirasawa and Feil, 2008) (Fig. S2B).

At the zygote stage, 856 transcripts were present (FPKM >10) in human and marmoset, but not in mouse (Fig. 2E, S2C). Pathway and Gene Ontology (GO) analyses identified genes associated with “ribosome”, “cytoplasmic ribosomal proteins” and “translational termination”, indicating an abundance of transcripts involved in translational processing (Fig. S2D). Consistent results were obtained with individual GO term analyses confined to each species (Fig. S2E–G). We compared enrichment scores of the most significant processes identified in human to those in marmoset and mouse (Fig. 2F). General biological processes, such as “gene expression” and “mitotic cell cycle” were elevated in all species. However, terms associated with translation were highly enriched in primates and lacking in mouse. We also assessed individual ribosomal transcript RNA levels and found substantial reductions in mouse relative to primates (Fig. 2G). These observations suggest that primate maternal programs are adapted to ensure extended translation of maternally deposited RNAs until later ZGA at the 8-cell stage.

Mouse maternal effect genes include *de novo* and maintenance DNA methyltransferases *Dnmt3a* (Okano et al., 1999) and *Dnmt1* (Howell et al., 2001). We examined chromatin remodelling factors by hierarchical clustering (Fig. 2H, Table S2). In marmoset and human, zygotes displayed higher levels of *TAF9*, *HDAC3* and *CTR9* transcripts. *DNMT1* was abundant in primates whereas *DNMT3A* and *DNMT3B* were also conserved in mouse (Fig. 2I). Human *DNMT3L* was present only at low levels in the zygote and 4-cell embryo, but elevated at the 8-cell stage and further upregulated in compacted morulae and early ICM; the marmoset followed a similar trend (Fig. 2I). This may suggest a requirement post-ZGA. We further observed that transcript levels of key members of the Polycomb Repressive Complex 1 and 2 (PRC1/2, Beisel and Paro, 2011; Morey et al., 2015), including *EED*, *SUZ12*, *EZH1* and *EZH2*, were substantially diminished or absent in primate zygotes (Fig. 2H). Strikingly, PRC components were upregulated in human and marmoset between 4- to 8-cell stages, corresponding to major ZGA timing (Fig. 2J). We conclude that PRC1 and PRC2 expression correlates with the onset of ZGA and is synchronous with species-specific developmental timing.

### Stage-specific transcription in rodent and primate embryos

We applied self-organising maps (SOM, Kohonen 1982) to define stage-specific gene expression modules. This allowed sorting of transcripts by expression pattern into 900 clusters per species, and identification of genes associated with each of the six developmental stages (Fig. 3A, S3A, C, Table S3). Statistical enrichment identified “sexual reproduction” and “chromosome segregation” as processes associated with zygotes of all three species (Fig. 3A, B, S3A–D). Human and marmoset 4-cell stages exhibited evidence of “negative regulation of transcription from RNA polymerase” (Fig. 3B, S3B), suggesting ZGA may be actively repressed in primates. The 8-cell stage in human and marmoset was dominated by terms relating to ZGA, while mouse cells showed enrichment for the JAK-STAT cascade (Fig. 3B, S3B, D). In late ICM we observed enrichment of “endodermal cell differentiation” and “extracellular matrix”, consistent with PrE segregation. Comparison of signalling pathways between species revealed “Phosphatidylinositol signalling” as conserved in zygotes, “Spliceosome”, “RNA transport” and “Basal transcription factors” at the 8-cell stage, and “Lysosome”, “Oxidative phosphorylation” in the early primate ICM (Fig. 3C–E).

**Fig. 3:**
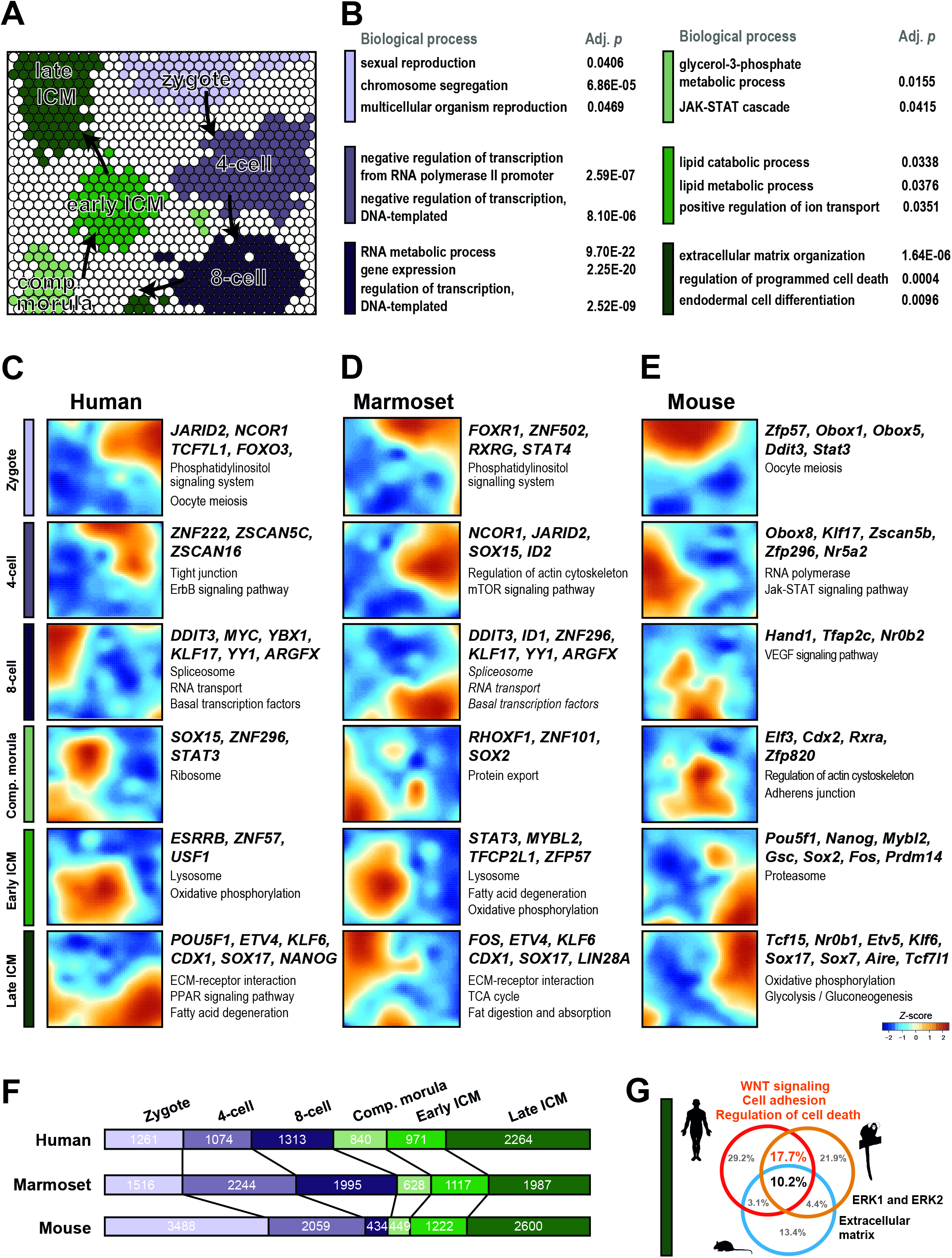
Stage-specific expression modules of preimplantation development. (A) Self-organizing map (SOM) of developmental stages from marmoset data. Stage-specific clusters (*Z*-score >1.5) are indicated by colour. (B) Enriched biological processes for specific SOM clusters. (C–E) SOM of human, marmoset and mouse stages, selected transcription factors and significantly enriched (*p* <0.05) KEGG pathways. (F) Numbers of stage-specific genes in each species. (G) Significantly enriched (*p* <0.05) biological processes at the late ICM stage.

We then extracted stage-specific transcription factors as candidates of regulatory interest (Fig. 3C–E). Human and marmoset shared *JARID2* prior to ZGA and concomitantly upregulated *DDIT3*, *KLF17* and *YY1*, for which deletions in mouse are embryonic lethal (Donohoe et al., 1999), at the 8-cell stage. The maternal effect gene *Zfp57*, for which roles in imprint maintenance have recently been characterised (Strogantsev et al., 2015; Riso et al., 2016), was specific to mouse zygotes (Fig. 3E) but subsequently expressed in the early marmoset ICM (Fig. 3D). Human *ZFP57* expression followed the pattern observed in marmoset (Table S3). In the late ICM, we found conserved expression of *ETV4*, *SMAD6*, *KLF6* and PrE markers *SOX17* and *FOXA2* in all species (Fig. 3C–E). Interestingly, the late mouse ICM alone expressed the pluripotency repressor *Tcf7l1* (*Tcf3*) (Wray et al., 2011; Yi et al., 2011) together with *Tcf15* and ETS-related factor *Etv5*, implicated in the onset of embryonic stem cell differentiation (Davies et al., 2013; Akagi et al., 2015) (Fig. 3E), The presence of such antagonists of the pluripotency network may contribute to accelerated progression of embryonic development in mouse relative to primates.

We next quantified the distribution of stage-specific genes between the three species (Fig. 3F). Human and marmoset both displayed substantially fewer zygote-specific transcripts (1261 and 1517, respectively) than mouse (3488). The inverse pattern was observed in the 8-cell embryo, correlating with ZGA in primates. By the end of the time series in late ICM cells, the numbers of stage-specific genes were similar in all species (Fig. 3F). This set of expression modules allowed us to define common and primate-specific processes at each developmental time point. Surprisingly, at the 8-cell, compacted morula and early ICM stages we found little correspondence between rodents and primates (Fig. S3E–G). In human and marmoset, the 8-cell transcriptome exhibited clear signs of ZGA, while the early ICM featured lipid metabolism. However, in the late blastocyst we identified several biological processes conserved in all species, including ERK signalling and upregulation of extracellular matrix components (Fig. 3G). We additionally identified a substantial fraction of primate-specific processes, primarily relating to WNT signalling, cell adhesion and apoptosis. This analysis faithfully captures known features of EPI and PrE segregation and identifies a number of distinct features associated with this lineage decision in late primate blastocysts.

### Non-human primate EPI and PrE specification

Cluster analysis indicated enrichment for both conserved and primate-specific features at EPI and PrE specification. This second lineage decision has been extensively described in mouse (Plusa et al., 2008; Artus et al., 2010; Artus et al., 2011; Artus and Hadjantonakis, 2012; Saiz and Plusa, 2013; Kang et al., 2013; Schrode et al., 2013; Schrode et al. 2014; Ohnishi et al., 2014), and more recently, human (Yan et al., 2013; Blakeley et al., 2015; Petropoulos et al., 2016; Stirparo et al., 2018), but remains poorly characterised in other species including non-human primates.

PCA of marmoset cells based on genome-wide expression resolved distinct sample groups by developmental time along the first dimension (Fig. 4A). Cells of the early ICM formed a cluster separate from compacted morulae and late ICM. Importantly, the late ICM began to diverge along the second dimension and, when those cells were examined in isolation, distinct EPI and PrE populations emerged (Fig. 4B, S4A). Pluripotency factors *TGDF1*, *NANOG*, *GDF3* and *KLF17* contributed to the EPI trajectory. Moreover, we found Activin/Nodal signalling components *NODAL* and *LEFTY2* prominent in the EPI cluster. Genes contributing to PrE segregation comprised *PDGFRA*, *LAMA1*, *APOE*, *SPARC* and *RSPO3*, a positive regulator of canonical WNT signalling (Nam et al., 2006).

**Fig. 4:**
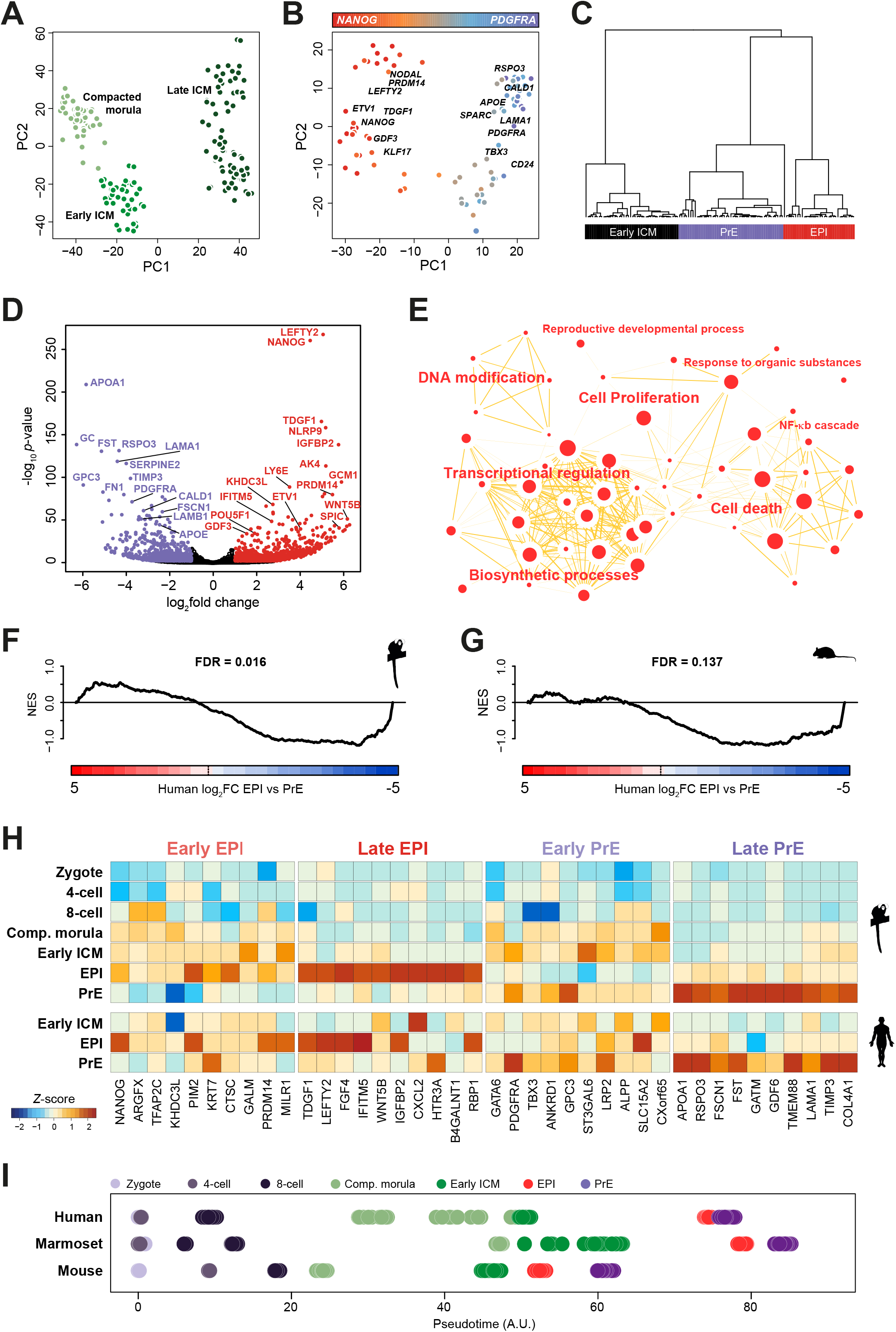
The second lineage decision in the marmoset. (A) PCA of marmoset samples from compacted morula, early and late ICM stages (FPKM >0). (B) PCA based on variable genes (log_2_FPKM >0, logCV^2^ >0.5, *n*=3363) for the marmoset late ICM. (C) Weighted gene co-expression network analysis (WGCNA) represented as clusters of eigengene values for early and late ICM. (D) Genes differentially expressed between marmoset EPI (red) and PrE (purple). (E) Cytoscape enrichment map of the top 50 biological processes (*p* >0.05) based on absolute fold change >0.5 between PrE and EPI. (F–G) Gene set enrichment analysis (GSEA) based on genes differentially expressed between (F) human and marmoset and (G) human and mouse EPI versus PrE. (H) Representative early and late EPI and PrE markers in marmoset and human. (I) Pseudotime analysis of human, marmoset and mouse embryonic lineages.

To independently assess whether early ICM, EPI and PrE cells represent distinct populations, we performed weighted gene co-expression network analysis (WGCNA, (Zhang and Horvath, 2005)) based on highly variable genes (Methods). We extracted three major co-expression modules by unsupervised clustering, corresponding to early ICM, PrE and EPI (Fig. 4C). These populations were consistent with cell fate assignment based on PCA (Fig. 4B, S4A).

We performed differential analysis between marmoset EPI and PrE expression signatures (Fig. 4D, Table S4). The EPI transcriptional network contained *NANOG, LEFTY2* and *TDGF1*, whereas we identified *APOA1*, *RSPO3*, *GPC3*, *FN1*, *PDGFRA* and *LAMA1* as PrE markers. Notable among the top EPI-specific genes was *WNT5A*, further suggesting a role for WNT signalling in lineage segregation. Differentially enriched biological processes in PrE featured cell migration, adhesion and lipid metabolism (Fig. S4B). EPI-enriched processes included transcriptional regulation, proliferation and a complex network of cell death associated nodes (Fig. 4E), consistent with our cross-species comparison (Fig. 3G). Marmoset EPI contained a “DNA modification” module (Fig. 4E), and the *de novo* DNA methyltransferase *DNMT3B* was among the top 25 differentially expressed genes in marmoset EPI versus PrE (Table S4).

We used gene set enrichment analysis (GSEA, Subramanian et al., 2005) to compare EPI versus PrE transcriptional signatures between species (Fig. 4F, G). There was significant concordance of genes differentially expressed between EPI and PrE in human and marmoset (Fig. 4F), but not human and mouse (Fig. 4G). Pearson correlation of marmoset EPI to human and mouse EPI was significantly higher in human, and similar results were obtained for PrE (data not shown). Collectively these results support conservation in transcriptional networks in late primate ICM.

Analysis of all marmoset preimplantation stages based on variable genes showed unambiguous segregation of EPI and PrE, even in the presence of pre-ZGA stages (Fig. S4C). To define robust marker sets for EPI and PrE lineage acquisition in primates, we derived developmental trajectories based on pseudotime. Plotting known pluripotency and germ-cell associated genes allowed us to discern expression levels, temporal dynamics, heterogeneity and EPI/PrE lineage association (Fig. S4D). We confirmed absence of mouse-specific pluripotency markers *FBXO15*, *GBX2*, *ESRRB*, *UTF1* and *KLF2* in the marmoset preimplantation epiblast, similar to recent reports in human (Blakeley et al., 2015; Stirparo et al., 2018). *NANOG*, *ARGFX, TFAP2C, MILR1* and *KHDC3L* were already expressed in morulae and early ICM and retained in the EPI, but downregulated in PrE (Fig. 4H). Conversely, *TDGF1, LEFTY2, FGF4*, *IFITM5* and *IGFBP2* were expressed at low abundance at earlier stages, but sharply upregulated upon EPI specification. We identified *GATA6*, *LRP2, TBX3* and *ANXA3* as early PrE markers robustly expressed prior to PrE specification, while *APOA1*, *COL4A1*, *GDF6*, *RSPO3* and *FST* were exclusive to the mature PrE lineage.

We then mapped individual transcriptomes on a temporal trajectory for each species. Stage- and lineage-specific groups were largely apparent, recapitulating the relative duration of preimplantation development (marmoset > human > mouse) and early embryo progression characteristic of each species (Fig. 4I). We conclude that marmoset cells readily segregate into discrete clusters of early ICM, EPI and PrE and mirror global features of human preimplantation development, including primate-specific marker acquisition and expression of canonical WNT signalling components. Temporal analysis further suggests that EPI is likely specified prior to PrE (Grabarek et al., 2012).

### Transposcriptome signatures of rodent and primate preimplantation stages

We sought to define stage-specific transposable element signatures of rodent and primate preimplantation development. Transcription was detected in human embryo cells from more than 100,000 of 4,000,000 annotated repeat loci. Analysis of variably expressed elements largely distinguished developmental stages in human (Fig. 5A), marmoset (Fig. 5B) and mouse (Fig. 5C), although such classification was less definitive than analyses based on gene expression (Fig. 1C–E). Nevertheless, PCA of late ICM cells based on transposable elements accurately segregated EPI and PrE populations in all three species (Fig. S5A–C).

**Fig. 5:**
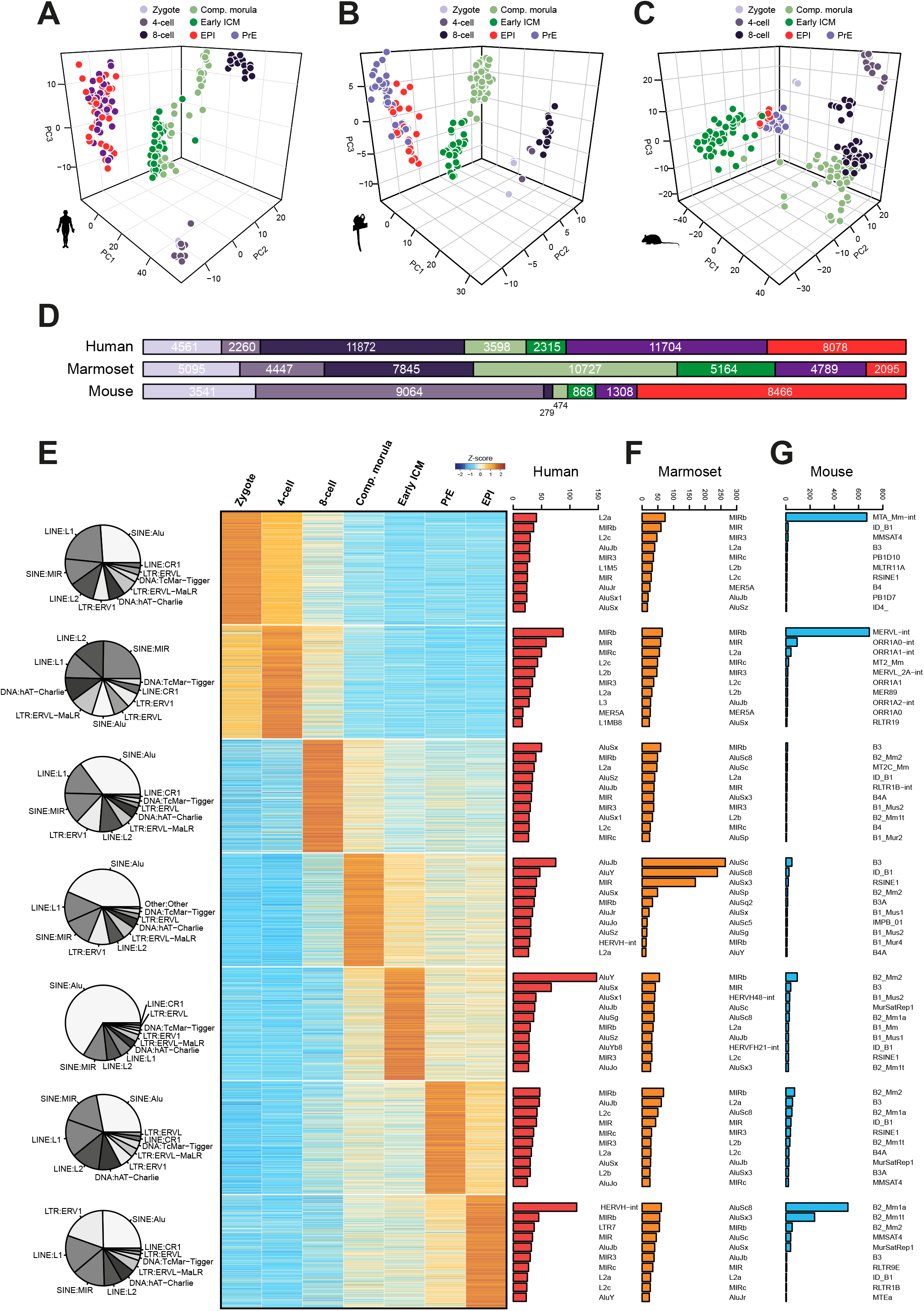
The transposcriptome in preimplantation development. (A–C) PCA of selected transposable elements (log_2_ normalised count >0.5 and logCV^2^ >1) expressed in human (A), marmoset (B) and mouse (C). (D) Numbers of stage-specific transposable elements for all preimplantation stages (for individual elements = *Z*-score >2 and normalised read counts >10). (E) Top 1000 stage-specific transcripts in human. Pie charts indicate proportions of the 10 most abundant classes for the top 1000 stage-specific transposable elements. Bar charts display counts for the 10 most abundant families encompassing the top 1000 stage-specific elements. (F–G) Most abundant retrotransposon families for the top 1000 stage-specific transcripts in marmoset (F) and mouse (G) as defined in Table S7.

To elucidate dynamics of transposon expression in human embryos we performed hierarchical clustering of 925 sequence families (Fig. S5D, Table S5). The majority were robustly expressed during early cleavage, with a substantial decline after the 8-cell stage. Similarly, marmoset cells showed extensive downregulation at the transition from 8-cell embryos to compacted morulae (Fig. S5E). In the mouse, a large fraction of transposable element families was upregulated at the 4-cell stage and the majority sustained robust expression beyond 8-cell (Fig. S5F). We also examined families associated with naïve versus primed pluripotency in cultured human pluripotent stem cells (PSC) (Theunissen et al., 2016; Collier et al., 2017), and found robust expression of SINE/VNTR/Alu (SVA) elements, specifically SVA_B, SVA_C, SVA_D, SVA_E, SVA_F, as well as HERVH-int and LTR5/7 in human EPI (Fig. S5G). However, other families reported to be associated with naïve (SVA_A, LTR49-int) and conventionally cultured (LTR7C, MSTA-int and THE1D-int) PSC were minimally expressed or undetectable in EPI. A significant proportion of transposable elements detected in naïve cultures exhibited higher expression at the 8-cell and compacted morula stages than EPI, consistent with previous findings (Theunissen et al., 2016).

Transposcriptome analysis based on family association is limited by considerable heterogeneity in expression of individual elements. This prompted us to determine stage-specific profiles based on individual loci, rather than family or class affiliation. Applying stringent selection criteria for specificity (*Z*-score >2) and excluding marginally expressed transcripts (normalised counts <10) revealed more than 40,000 stage-specific elements in human and marmoset (Fig. 5D, Table S6). Mouse samples featured many transcripts enriched in zygote and 4-cell stages as well as the EPI lineage, but surprisingly few specific to eight-cell embryos (279), compacted morulae (474) and early ICM (868). This contrasted with a more uniform distribution in primates, where at least 2000 elements could be associated with any given stage.

To attempt more precise developmental staging on transposable element expression, we extracted the top 1000 (or the maximum available for three time points in mouse) specific to embryonic stages, including EPI and PrE, and assessed repeat class and family association in human (Fig. 5E), marmoset (Fig. 5F) and mouse (Fig. 5G). The most abundant classes in human were SINE:Alu, LINE:L1, LINE:L2 and SINE:MIR. The early ICM was characterised by pronounced expression of Alu families, including AluY, AluSx, AluSx1, AluJb and AluSg, and we found strong enrichment for HERVH-int, MIR2b and LTR7 in the EPI (Fig. 5E). Notably, SVAs did not feature in the top 10 transcript families of any preimplantation stage. Marmoset embryonic cells displayed similarly high abundance of primate-specific Alu families, in particular AluSc, AluSc8 and AluSx3 (Fig. 5F). The mouse expression profile differed profoundly from primates and revealed strong specificity for MTA_Mm-int in zygote, MERVL-int at the 4-cell stage and B2_Mm1a, B2_Mm1t and B2_Mm2 in the EPI (Fig. 5G). In summary, these results suggest that transposable elements can be used to discern early stages of embryonic development based on the repertoire of sequences expressed. We note considerable differences in transposable element composition between rodents and primates, and were able to resolve expression signatures for distinct preimplantation embryo lineages.

### Conserved and primate-specific elements of the EPI and PrE transcription factor networks

We sought to derive transcription factor networks and extract a set of conserved regulators of primate pluripotency in the EPI lineage (Fig. 6A). More transcription factors were shared between human EPI and marmoset (139) than mouse (47). We constructed a core pluripotency network common to all species, based on the 282 transcription factors shared in the EPI and excluding those expressed in PrE (Fig. S6A). The conserved EPI network included the core pluripotency factors *POU5F1*, *SOX2* and *NANOG* as well as *TFCP2L1*, *ZNF296*, *ZFP36L1* and *SOX15*. We also noted the presence of transcriptional repressor *RBPJ* and ERK-signalling associated factors *ETV4* and *ETV5*.

**Fig. 6:**
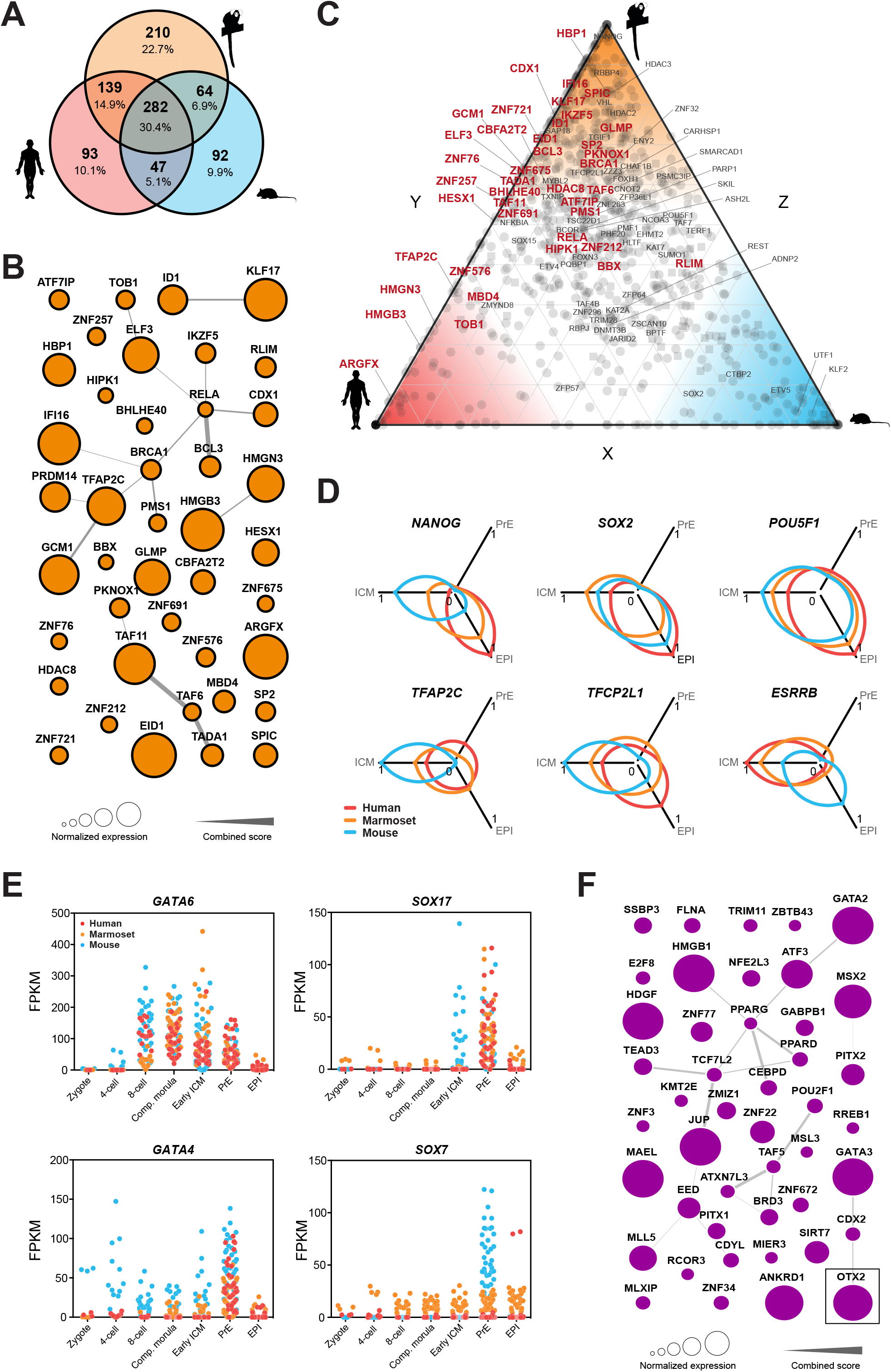
Conserved and divergent elements of EPI and PrE transcription factor networks. (A) Intersection of transcription factors specific to EPI (FPKM >5 in EPI and not significantly (*p* >0.05) upregulated in PrE). (B) Protein-protein interaction network of primate-specific EPI transcription factors. Node sizes are scaled to normalised expression in human and marmoset; edges are derived from the STRING database. (C) EPI enriched transcription factors (circles) and chromatin remodelling factors (squares). Axes show the relative fraction of expression in the EPI between mouse and human (*x*), human and marmoset (*y*), and marmoset and mouse (*z*). (D) Selected markers representing normalised expression in ICM, EPI and PrE. (E) Sequentially activated canonical mouse PrE markers expressed in mouse (blue), marmoset (orange) and human (red). (F) Protein-protein interaction network of primate-specific PrE transcription factors (FPKM >5 in PrE and not significantly (*p* >0.05) upregulated in EPI). As in (B), node sizes are scaled to normalised expression in human and marmoset and edges are derived from the STRING database.

We next assembled a network based on EPI-specific regulatory genes expressed in human and marmoset, but not mouse (Fig. 6B). We identified a substantial fraction of transcriptional repressors (*IKZF5*, *IFI16*, *BHLME40* and *CBFA2T2*), genes associated with DNA mismatch repair (*PMS1*, *MBD4*) and several components of NFκβ (*BCL3*, *RELA*) and WNT (*CDX1*, *HBP1*, *HMGN3*) signalling pathways. Interestingly, the network also contained *TFAP2C* and *GCM1*, which in mouse are associated with trophoblast lineage specification (Kaiser et al., 2015; Sharma et al., 2016) and regulation of syncytium formation (Kashif et al., 2011; Lu et al., 2016), respectively. For quantitative visualisation of species specificity we plotted relative expression of transcription factors and epigenetic modifiers (Fig. 6C). Transcription factors exclusively expressed in mouse EPI were *Klf2*, *Tcf15* and *Cited1*. Human EPI exhibited the highest levels of *SOX4* and *HAND1*, while *NANOG* expression was substantially elevated in marmoset.

We examined the dynamics of selected pluripotency factor expression in each species (Fig. 6D). Although *POU5F1*, *SOX2* and *NANOG* transcript levels correlated in equivalent lineages, *Nanog* was upregulated earlier in mouse. *TFAP2C* and *TFCP2L1* followed this pattern, whereas *ESRRB* was expressed substantially earlier in primate embryos and subsequently downregulated in EPI. We also found differences between primate species. Human-specific EPI factors included *CREB3L1*, *VENTX*, *HEY2* and *INSR*. Variations in insulin receptor (*INSR*) expression are potentially relevant to further refine human PSC culture conditions.

The conserved core transcription factor network for human, marmoset and mouse PrE contained many established lineage markers, including *GATA6*, *SOX17*, *HNF4A* and *GATA4* (Fig. S6B). We identified pluripotency-associated factors *TBX3* and *KLF5*, downstream targets of BMP signalling (*ID2*, *ID3*) and new potential regulators (*WDR77*, *PDLIM1*, *PCBD1* and *GTF2A2*) within the common PrE circuitry. In mouse, PrE lineage specification occurs via sequential activation of *Gata6* > *Sox17* > *Gata4* > *Sox7* (Artus et al., 2011). Consistent with this model we found that *Gata6* was expressed from the 8-cell stage, *Sox17* and *Gata4* were upregulated from early ICM formation and *Sox7* expression commenced in mature PrE (Fig. 6E). Human and marmoset *GATA6*, *SOX17* and *GATA4* largely followed this pattern, suggesting a potentially similar regulatory cascade underlying PrE specification in primates. However, *SOX7* remained low in marmoset throughout all developmental stages and absent in human.

To identify primate-specific regulators associated with PrE specification, we selected transcription factors expressed in human and marmoset PrE and excluded those in mouse PrE (Fig. S6C, D). We noted pronounced expression of the DNA chaperone *HMGB1*, regulator of desmosomes and intermediate junctions *JUP*, and endothelial transcription factors including *ANKRD1*, *GATA2* and *GATA3*. Surprisingly, the mouse ICM and postimplantation EPI-associated gene *OTX2* (Buecker et al., 2014; Acampora et al., 2016) was robustly expressed in the primate PrE (Fig. 6F).

In mouse, *Otx2* is first expressed in the ICM, then the preimplantation EPI (Acampora et al., 2016) and is subsequently highly upregulated upon implantation (Boroviak and Nichols, 2017). We implemented a resource to catalog and visualise embryonic gene expression in the species analysed (app.stemcells.cam.ac.uk/GRAPPA), and compared *OTX2* patterns in each time series. We observed a consistent pattern for mouse in our dataset (Fig. 7A, Table S7). In human and marmoset, however, *OTX2* is a maternal factor and specifically upregulated in PrE (Fig. 7A, Table S7), suggesting primate-specific adaptations of conserved transcriptional regulators for EPI and PrE segregation.

**Fig. 7:**
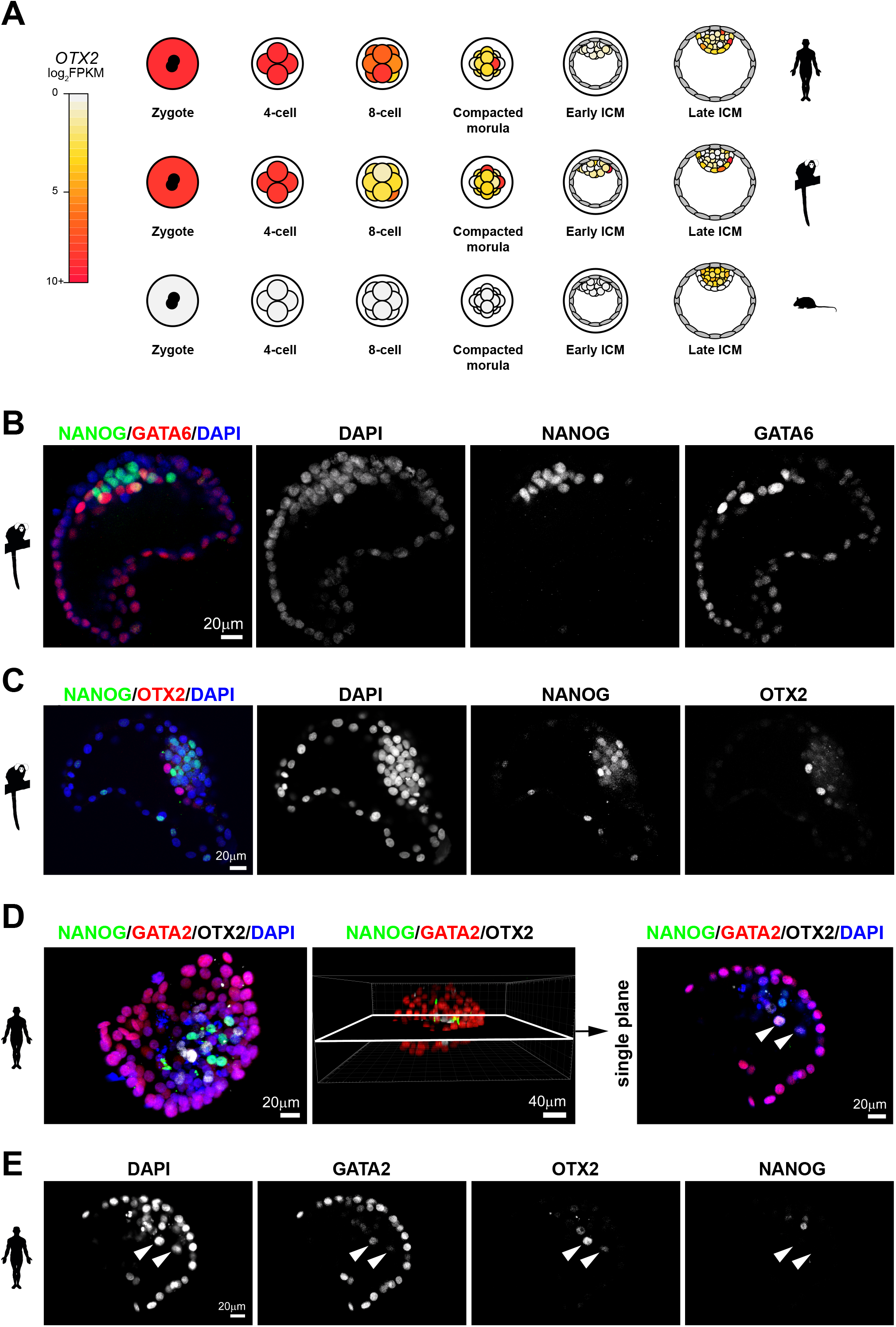
OTX2 protein localisation in primate embryos. (A) Schematic of Otx2 expression over preimplantation development. (B,C) Confocal microscopy immunofluorescence images of (B) NANOG, GATA6 and DAPI, and (C) NANOG, OTX2 and DAPI in marmoset late blastocysts. (D) Confocal sections, 3D reconstruction and single-plane image of NANOG, GATA2, OTX2 and DAPI localisation in an early human blastocyst. (E) Confocal sections of the indicated markers in a representative late human blastocyst. White arrows indicate PrE cells.

To determine whether *OTX2* transcript expression in primate PrE was reflected at the protein level we performed immunostaining. Marmoset embryos at this stage had completed EPI and PrE segregation as evidenced by mutually exclusive detection of NANOG and GATA6 (Fig. 7B). OTX2 was expressed in a subset of cells overlying the NANOG-positive EPI inside the blastocyst cavity (Fig. 7C). In human blastocysts at embryonic day (E)7 we also observed mutually exclusive staining between NANOG and OTX2 (Fig. 7D). We found GATA2 was absent in EPI, present at intermediate levels in PrE and highly expressed in all trophectoderm cells (Fig. 7E). OTX2 colocalised with intermediate levels of GATA2 inside the blastocyst, corroborating PrE-specific expression.

Rodent Otx2 directly binds regulatory genomic sequences of *Nanog* (Acampora et al., 2016). We therefore investigated the relationship of *NANOG* and *OTX2* transcriptional activity in each species at the single-cell level. *Nanog* and *Otx2* were co-expressed in all cells of the mouse EPI (Fig. S7A), whereas in human and marmoset *OTX2* was absent in half of EPI cells (marmoset 20/32, human 20/54) and expressed at low levels in the remainder (Fig. S7B, C). Mouse *Otx2* was consistently absent in PrE cells (41/44) (Fig. S7D). In contrast, *OTX2* was expressed in the majority of primate PrE cells (marmoset 34/47, human 32/38) (Fig. S7E, F). In the human embryo, cells with the highest levels of *OTX2* were devoid of *NANOG*.

This led us to investigate the dynamics of OTX2 and NANOG proteins at an earlier time point. We found that in cavitating human blastocysts (<100 cells) NANOG and OTX2 were co-expressed at intermediate abundance (Fig. S7G). However, in cells with strong signal for either NANOG or OTX2, expression was mutually exclusive. Similar patterns were observed in fully cavitated embryos (>100 cells) (Fig. S7H). These results may indicate a role for OTX2 in the regulation of EPI versus PrE lineage commitment in the human embryo, and the potentially divergent function of an established rodent pluripotency regulator in early primate development.

## Discussion

This study assembles a compendium of single-cell transcriptional data from preimplantation embryos of three mammalian species, and defines the regulatory events leading to rodent and primate EPI and PrE lineage specification. Features distinct to human and marmoset development are evident from the zygote stage. The primate maternal program is substantially enriched for ribosome biogenesis and components of the translational machinery. We identified a number of transcripts present in human and marmoset zygotes that are later confined to the EPI. Interestingly, *OTX2*, a homeobox transcription factor essential for mouse anterior forebrain identity (Perea-Gomez et al., 2001) and implicated in the progression of pluripotency (Acampora et al., 2013; Buecker et al., 2014; Yang et al., 2014; Kalkan et al., 2017), was present at high levels in both human and marmoset zygotes and later expressed in PrE.

We also identified notable differences with regard to epigenetic modifiers. The maintenance DNA methyltransferase *DNMT1* was highly upregulated in primates. In mouse *Dnmt3l* is required for the establishment of maternal imprints (Bourc’his et al., 2001), but *DNMT3L* was completely absent in human and marmoset. Core members of PRC1 and PRC2 complexes, including *BMI*, *EED*, *EZH1* and *SUZ12* were scarcely detectable in the primate zygote, but upregulated between 4- and 8-cell stages. In mouse, PRC1/2 are already established in the zygote. The presence of key PRC1/2 components at ZGA suggests a role for repressive complexes immediately following engagement of the transcriptional apparatus.

The marmoset early ICM embodies a distinct transcriptional state, in agreement with recent reports in human (Blakeley et al., 2015; Stirparo et al., 2018), *Macaca fascicularis* (Nakamura et al., 2016) and mouse (Mohammed et al., 2017). Primate early ICM cells expressed essential regulators of early mouse embryonic development, including *POU5F1*, *GATA6*, *STAT3* and *MYBL2*. Transcripts exclusive to primate ICM were *NUCB1*, encoding a calcium binding protein of the Golgi, *STEAP1*, a metalloreductase, and *DLC1*, a GTPase-activating protein involved in cytoskeletal reorganisation and activator of phospholipase PLCD1 which also peaks during ICM formation. Cluster analysis revealed enrichment for lipid metabolism in human and marmoset early ICM, suggesting distinct metabolic requirements.

Progression from the early ICM leads to the establishment of naïve pluripotency in the EPI. Most current knowledge of the emergence and maintenance of pluripotent cells is derived from studies in mouse. We (Boroviak et al., 2015; Stirparo et al., 2018) and others (Blakeley et al., 2015; Nakamura et al., 2016) demonstrated that a substantial fraction of mouse pluripotency-associated factors are absent from human and non-human primate ICM, including *KLF2*, *NR0B1*, *ESRRB*, *FBXO15* and *JAM2*. Cross-species analyses defined primate-specific transcription and chromatin remodelling factor circuitry in the EPI. We identified *TFAP2C* and *GCM1*, regarded as TE-associated factors in mouse, in the naïve primate pluripotency network with *KLF17*, *ARGFX*, *HESX1*, *ELF3, HMGB3* and several zinc-finger proteins, including *ZNF675*, *ZNF257*, and *ZNF146*. It is tempting to speculate that a subset of these regulators may endow primate EPI cells with the potential for amnion specification directly after implantation (Boroviak and Nichols, 2017).

Comparative analyses of rodent and primate EPI defines conserved core pluripotency factors, in addition to *POU5F1*, *SOX2* and *NANOG*. These include *TFCP2L1* (Martello et al., 2013; Ye et al., 2013), *ZSCAN10*, shown to promote genomic stability in mouse embryonic stem cells (Skamagki et al., 2017), *ZNF296*, a component of heterochromatin (Matsuura et al., 2017) reported to enhance reprogramming efficiency (Fischedick et al., 2012) and interact with *KLF4* (Fujii et al., 2013), *ZFP36L1*, a zinc-finger RNA-binding protein attenuating protein synthesis (Stoecklin et al., 2002) and *MYBL2*, which regulates cell cycle progression and is essential for mouse ICM formation (Tanaka et al., 1999). The conserved pluripotency network further contained repressive chromatin remodelling factors *HDAC2*, *HDAC3* and *JARID2* as well as ETS-related transcription factors (*ETV4*/*5*), involved in cell proliferation and induction of differentiation-associated genes in mouse embryonic stem cells (Akagi et al., 2015). Interestingly, we also found *RBPJ*, a transcriptional repressor regulated by Notch signalling, to be an integral part of the EPI program in all three species despite the absence of Delta/Notch receptors. Mouse *Rbpj* is not required for self-renewal in embryonic stem cell lines, but is implicated in early differentiation and neural lineage entry (Lowell et al., 2006; Leeb et al., 2014).

Repetitive DNA sequences have been introduced into mammalian genomes over evolutionary time, and constitute approximately half of the human genome. Transposable elements are subject to epigenetic control and expression during early embryogenesis, and associated transcription may provide a signature to define developmental states. Correspondence in transposcriptome has been proposed as a metric for comparison of cultured human PSC lines to *in vivo* counterparts (Theunissen et al., 2016; Collier et al., 2017). We derived transposable element-based signatures for six embryonic stages, including EPI and PrE, and noted considerable differences in transcript repertoire between species. Transposable element families reported in human PSC were largely expressed in the EPI. Despite that general finding, SVA_A and LTR49-int were barely detectable in the EPI, and instead expressed at earlier stages. Similarities in transposcriptome profiles of PSC cultures and morula cells may be a consequence of hypomethylation *in vitro* (Theunissen et al., 2016; Pastor et al., 2016).

Acquisition of PrE identity in primates largely recapitulated the sequential activation of lineage specifiers in mouse. *GATA6* was robustly expressed in morulae and early ICM and subsequently confined to PrE in the late ICM, while *GATA4* and *SOX17* were specifically upregulated in PrE. We determined *SOX7* to be absent in primates. Notably, mouse Sox7 is dispensable for PrE formation from mouse embryonic stem cells (Kinoshita et al., 2015), in contrast to the potent inductive function of *GATA6* (Fujikura et al., 2002), *SOX17* (McDonald et al., 2014) and *GATA4* (Fujikura et al., 2002).

We identified an array of new core PrE-associated factors present in human, marmoset and mouse. These included *KLF5*, *KLF6*, *TBX3*, *EGR1* and BMP signalling components *ID2*/*3*. Cluster analysis revealed enrichment for ERK signalling in all three species. However, we identified the WNT pathway as a primate-specific feature in the late ICM. Primate PrE consistently expressed *RSPO3*, a potent WNT signalling enhancer (de Lau et al., 2014). Human EPI featured *WNT3*, (marmoset *WNT5B*) in contrast to complete absence of WNT ligand in mouse. We have previously shown that WNT inhibition interferes with NANOG and GATA6 segregation in the marmoset embryo (Boroviak et al., 2015), supporting a functional requirement for WNT in primate PrE specification.

The primate-specific PrE transcription factor network contained *ANKRD1*, *PITX2*, *MSX2*, *CEBPD*, *HDGF* and mouse trophoblast markers *GATA2* and *GATA3*, which were robustly expressed in PrE, although orders of magnitude lower than in the TE lineage. A previous report has proposed *DPPA4* as a new human PrE marker (Petropoulos et al., 2016). However, our re-analysis of human datasets (Stirparo et al., 2018) with stage-matched marmoset and mouse samples showed that *DPPA4* is consistently upregulated in the EPI. Mouse-specific PrE factors were *Tfec*, *Sox7*, *Foxq1*, *Cited1* and *Hhex*. The homeobox gene *Hhex* is first expressed in PrE and during implantation confined to the distal tip of visceral endoderm (Thomas et al., 1998), which becomes the anterior visceral endoderm. Mouse *Hhex*-expressing PrE cells have a tall, columnar epithelial morphology, overlying the EPI in a distal position (Rivera-Perez et al., 2003). The absence of *HHEX* in human and marmoset PrE suggests potentially divergent mechanisms for the establishment of asymmetry in primates. Strikingly, we discovered *OTX2* as a PrE associated gene in primates. In mouse, Otx2 is not present in PrE but is later required for visceral endoderm movement and for the restriction of posterior signals in the EPI (Perea-Gomez et al., 2001). We show that in primates OTX2 protein localises in a subset of PrE cells and becomes mutually exclusive with NANOG at the late blastocyst stage. Further studies are required to determine the role of OTX2 in primate PrE formation.

Collectively, we present a framework for comparative molecular evaluation of human embryology to a tractable non-human primate model and the established mouse paradigm. Our cross-species analysis resolves genome-wide expression signatures of protein-coding genes and transposable elements for major preimplantation embryo stages as a benchmark for *in vitro* cultured cells. This work provides a resource for high-confidence, primate-specific lineage factors for future functional interrogation.

## Materials and Methods

### Marmoset colony maintenance and embryo collection

Marmoset embryos were obtained from the Central Institute for Experimental Animals, Kanagawa, Japan (CIEA). Experiments using marmosets at the CIEA were approved by the animal research committee (CIEA: 11028) and performed in compliance with guidelines set forth by the Science Council of Japan. Marmosets were maintained as previously described (Hanazawa et al., 2012). Embryos were collected according to established methods using recently developed devices (Takahashi et al., 2014; Thomson et al., 1994). Staging of female marmoset reproductive cycles and embryo collection have been described (Hanazawa et al., 2012).

### Single-cell RNA-seq datasets

We compiled three stage-matched, single-cell preimplantation embryo datasets based on published and newly generated samples for human, marmoset and mouse. All samples were processed with the Smart-seq library construction method for full-length coverage of individual transcripts (Picelli et al., 2013). This resulted in a compendium of seven embryonic lineages from six developmental stages, spanning zygote, 4-cell, 8-cell, compacted morula, early and late ICM.

Human embryo transcriptomes were compiled from three single-cell profiling studies (Yan et al., 2013), (Blakeley et al., 2015), (Petropoulos et al., 2016). These data were processed and annotated based on the analysis reported in (Stirparo et al., 2018).

Marmoset samples were newly generated for this study. Embryos were staged according to cell number and embryonic day. Where appropriate, zona pellucidae were removed using acid Tyrode’s solution (Sigma) and embryos were subjected to immunosurgery as previously described (Solter and Knowles, 1975; Nichols et al., 1998) using a custom rabbit polyclonal anti-marmoset antibody (Boroviak et al., 2015). Following the complement reaction, residual trophectoderm was thoroughly removed by repetitive manual pipetting. For dissociation of marmoset ICM into single cells, recovered ICM were exposed to a 1:1 mixture of 0.025% trypsin plus EDTA (Invitrogen) and 0.025% trypsin (Invitrogen) plus 1% chick serum (Sigma) for 5–10min. Cells were dissociated into singletons by repetitive pipetting with micro-capillaries of gradually reduced inner diameter. Individual ICM cells were transferred to single-cell lysis buffer and snap frozen on dry ice. Smart-seq2 libraries were prepared as described (Picelli et al., 2014) and sequenced on the Illumina platform in 125bp paired-end format. These data are available via ArrayExpress accession E-MTAB-7078.

Mouse embryo data were compiled from an earlier study (Deng et al., 2014) and augmented by samples produced in our laboratories (Mohammed et al., 2017). Single-cell transcriptome profiling of mouse preimplantation embryos has been reported (Deng et al., 2014); however, that study did not yield samples at the late ICM stage representing distinct EPI and PrE lineages (Blakeley et al., 2015). We therefore produced stage-matched single-cell samples for early and late blastocyst ICM, and sequenced RNA-seq libraries prepared with the Smart-seq2 protocol. These data have been previously described (Mohammed et al., 2017).

### RNA-seq data processing

Sequencing data from single-cell human (accessions SRP011546 (Yan et al., 2013), SRP055810 (Blakeley et al., 2015), ERP012552 (Petropoulos et al., 2016)), and mouse (SRP110669 (Mohammed et al., 2017), SRP020490 (Deng et al., 2014)) embryo profiling studies were obtained from the European Nucleotide Archive (Silvester et al., 2018). Reads from each species dataset were aligned to human genome build GRCh38/hg38, common marmoset C_jacchus3.2.1, and mouse assembly GRCm38/mm10 with *STAR* 2.5.2b (Dobin et al., 2013) using the two-pass method for novel splice detection (Engström et al., 2013). Read alignment was guided by gene annotation from Ensembl release 87 and splice junction donor/acceptor overlap settings were tailored to the read length of each dataset. Alignments to gene loci were quantified with *htseq-count* (Anders et al., 2015) based on annotation from Ensembl 87 (Yates et al., 2016). Sequencing libraries with fewer than 500K mapped reads were excluded from subsequent analyses. Read distribution bias across gene bodies was computed as the ratio between the total reads spanning the 50th to the 100th percentile of gene length, and those between the first and 49th. Samples with ratio >2 were not considered further.

### Transcriptome analysis

Principal component and cluster analyses were performed based on log_2_ FPKM values computed with the Bioconductor packages *DESeq2* (Love et al., 2014), *Sincell* (Juliá et al., 2015) or *FactoMineR* (Lê et al., 2008) in conjunction with custom scripts. If not otherwise indicated default parameters were used. Differential expression analysis was performed with *scde* (Kharchenko et al., 2014), which fits individual error models for the assessment of differential expression between sample groups. For global analyses, genes that registered zero counts in all single-cell samples in a given comparison were omitted. Euclidean distance and average agglomeration methods were used for cluster analyses. Volcano plots were computed with log_2_ fold change and -log_10_ *p*-values from *DESeq2* differential analysis output. Mutual information scores were computed with the *infotheo* R package. Orthology mapping was performed according to gene annotation in Ensembl release 87 with human as the reference species. Multiple orthologies were deconvoluted based on the percentage of gene sequence similarity as defined in Ensembl. Expression data are available in supplementary Tables S2 and S4, and through a web application to visualise expression levels of individual genes in embryonic lineages (app.stemcells.cam.ac.uk/GRAPPA).

### Selection of high-variability genes

Genes exhibiting the greatest expression variability (and thus contributing substantial discriminatory power) were identified by fitting a non-linear regression curve between average log_2_ FPKM and the square of coefficient of variation. Thresholds were applied along the *x*-axis (average log_2_ FPKM) and *y*-axis (log CV^2^) to identify the most variable genes.

### Evaluation of refined embryonic cell populations

To assess the accuracy of selected EPI, PrE and early ICM cells, we used the Weighted Gene Co-Expression Network Analysis unsupervised clustering method (WGCNA, Langfelder and Horvath, 2008) to identify specific modules of co-expressed genes in each developmental lineage. A soft power threshold of 10 was set to govern the correlation metric and a tree pruning approach (Langfelder et al., 2008) was implemented to merge similar modules (threshold 0.35). The minimum module size was set to 50 genes; from the modules computed, the top 50 genes with greatest intramodular connectivity were selected for subsequent co-expression network analysis.

### Pseudotime analysis

Temporal trajectories were computed with the *monocle* R package (Trapnell et al., 2014), using the DDRtree reduction and vstExprs normalisation options. As different numbers of cells were profiled from each species and developmental stages in the datasets considered, uniform cell numbers were sampled from each group and the average was reported from 100 sampling iterations.

### Network analysis of biological processes

Statistical enrichment of Gene Ontology (GO) terms was computed with the *GOstats* Bioconductor package, DAVID 6.8 (Huang et al., 2009) and EnrichR web tools (Kuleshov et al., 2016). *Cytoscape* (Shannon et al., 2003; Smoot et al., 2011) and the associated enrichment map plugin were used for network construction and visualisation. For network diagrams, node size is scaled by the number of genes contributing to over-representation of biological processes; edges are plotted in widths proportional to the overlap between gene sets.

### Derivation of stage-specific expression modules

The R package *kohonen* was used to construct self-organising maps (SOM) across embryonic stages for human, marmoset and mouse. Variation in transcriptional activity was identified using a matrix of 30×30 with hexagonal topology. Stage-specific GO analyses were performed with *GOstats* package considering genes with *Z*-score >1.5, while genes with *Z*-score <1.5 in all stages were used for the background set. Annotation related to transcription factors, co-factors and chromatin remodelling factors was obtained from AnimalTFDB 2.0 (Zhang et al., 2015). Marmoset late lineage markers were selected as genes expressed in EPI or PrE cells with a transcriptional contribution more than 75% across all selected pre-implantation stages and minimum level of 10 FPKM. Early markers were identified as genes in later stages (from 8-cell morulae to either EPI or PrE lineages) with a transcriptional contribution of more than 75% across all selected pre-implantation stages. A fold change induction of at least four between lineages and minimum level of 10 FPKM in at least in one of the following stages was required: 8-cell, compacted morula, early ICM, EPI or PrE.

### Identification of EPI- and PrE-associated transcription factors

A two-step process was used to determine sets of transcription factors, co-factors and chromatin remodellers enriched in embryonic lineages of the species analysed. Genes expressed at greater than 5 FPKM in the subject lineage (e.g. EPI) and not significantly downregulated in the other (e.g. PrE) were selected in human, marmoset and mouse and averaged for all cells annotated in each cell type. Genes in common to all species, or alternatively were specific to primate, were compared between EPI and PrE modules. Ternary plots were produced with the R package *ggtern* using the relative percentage of average expression for all cells in EPI and PrE lineages. Protein-protein interactions between factors expressed in a primate-specific context or common to all species were computed based on entries curated in the STRING database (Szklarczyk et al., 2017).

### Transposable element analysis

RepeatMasker annotations for human, marmoset and mouse genomes were obtained from the UCSC Table Browser (Karolchik et al., 2004). To calculate expression levels for transposable elements, adapter-trimmed RNA-seq reads were aligned to the respective reference genome with *bowtie* (Langmead and Salzberg, 2012) using parameters ’-m1 –v2 --best --strata’ and selecting reads with unique alignment to individual elements, allowing two mismatches. Read counts for repeat regions and Ensembl transcripts were obtained by *featureCounts* (Liao et al., 2014), normalised by the total number of reads that mapped to Ensembl protein-coding transcripts, and subsequently normalised by repeat length. Differential expression between stages was evaluated with *DESeq2*.

### Immunofluorescence staining

Human and marmoset embryos were stained as previously described (Nichols et al., 2009; Boroviak et al., 2015). Primary antibodies were NANOG (Cell Signaling Technology 4893 (dilution 1:400)), GATA6 (R&D Systems AF1700 (1:100)), OTX2 (R&D Systems AF1979 (1:200)) and GATA2 (Abcam ab173817 (1:100)).

### Confocal imaging and analysis

Confocal images were acquired using a Leica TCS SP5 microscope. Optical section thickness ranged from 1–3 μm. Images were processed using Leica software, Imaris, Volocity and ImageJ (Fiji). Automated image analysis was performed in Volocity. Parameters for object identification were: guide size, 500μm^3^; separate objects, 500 μm^3^; and objects larger than 500μm^3^ were excluded. Thresholds for background fluorescence intensity (Gata6:45, Nanog:25, Cdx2:35, DAPI:20) were empirically determined to recapitulate manual cell counts in DMSO control embryos (Fig. S7A–D). For two IWP2-treated embryos the Nanog threshold was increased to 50 due to very bright signal to ensure accurate quantification of cell nuclei.

## Data availability

Single-cell RNA-seq data from marmoset embryos are available in the ArrayExpress repository under accession E-MTAB-7078.

## Funding

This work was supported by grants from the Biotechnology and Biological Sciences Research Council (BBSRC) UK (BB/M004023/1 (RG74277)), the Medical Research Council (MRC) UK (G1001028), and funding to the Cambridge Stem Cell Institute from the MRC and Wellcome Trust (097922/Z/11/Z, 203151/Z/16/Z). TB is a Wellcome Trust Sir Henry Dale Fellow. AS is an MRC Professor.

## Supplementary Figures

**Fig. S1:**
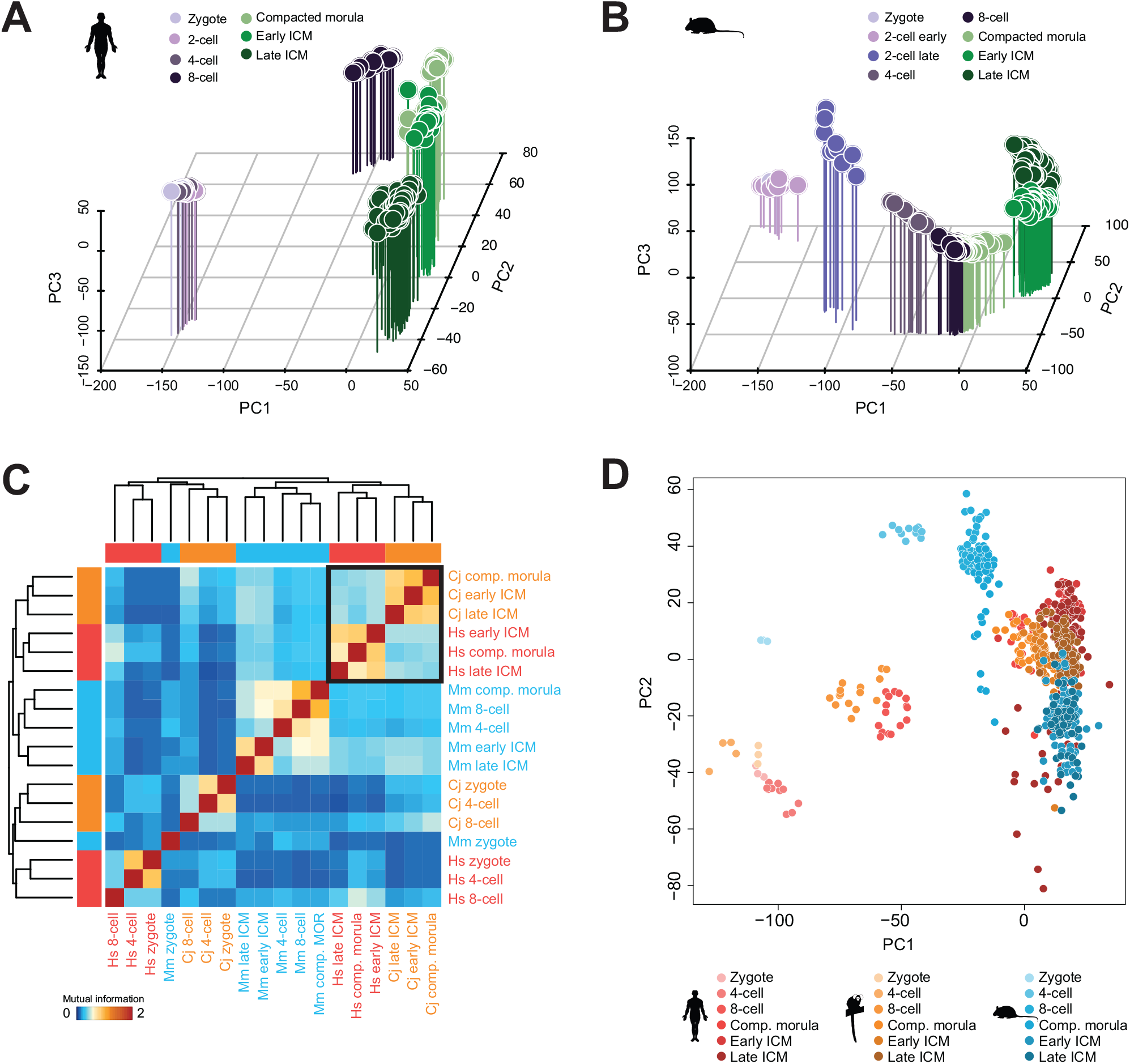
(A,B) PCA of single-cell embryo samples based on genes detected from minimal transcript coverage (FPKM >0) in human (A) and mouse (B). (C) Clustering of mutual information entropy for orthologous genes co-expressed in human, marmoset and mouse. (D) PCA of samples from all species based on orthologue *Z*-scores.

**Fig. S2:**
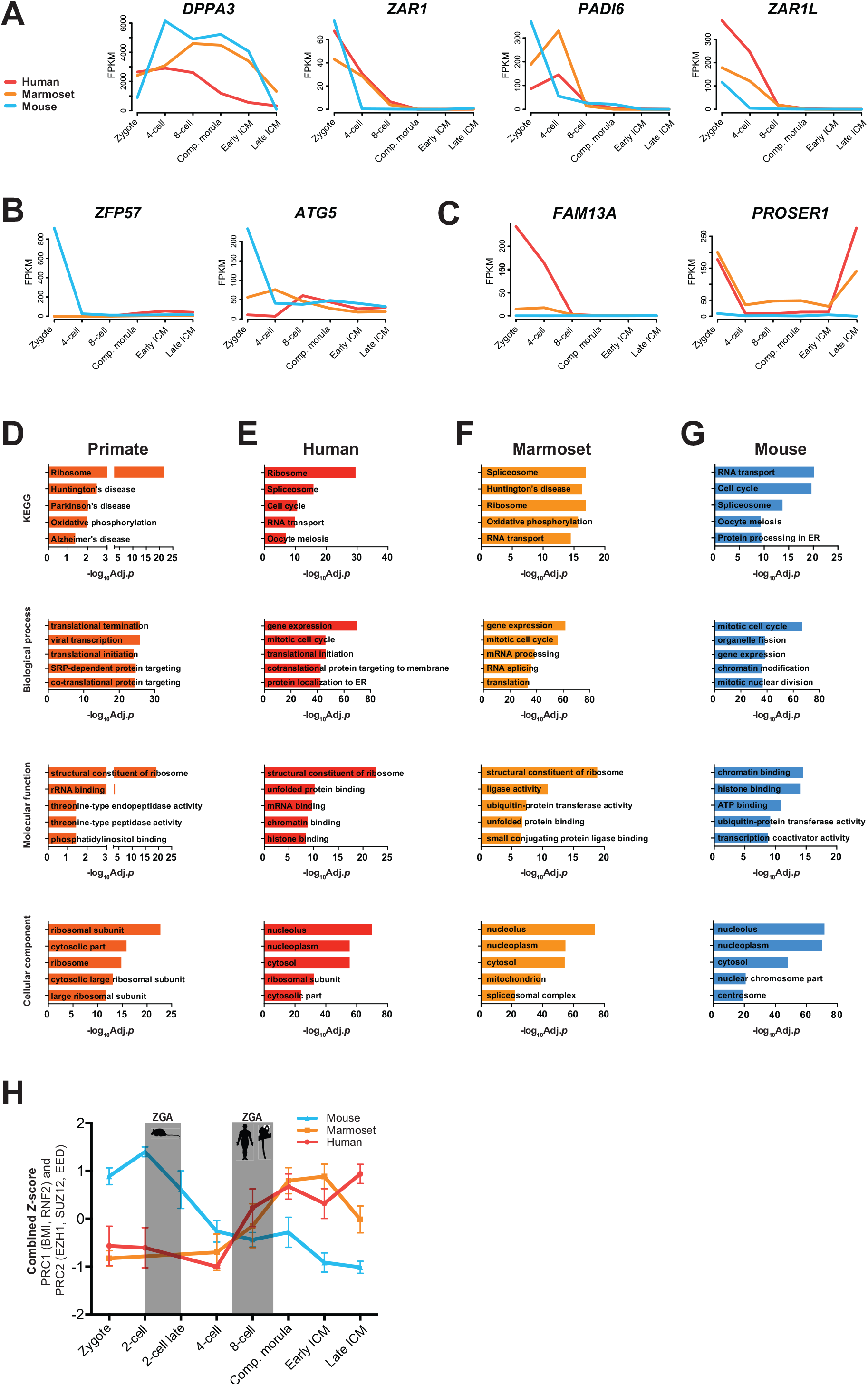
(A) Transcript abundance of conserved maternal effect genes in human, marmoset and mouse. (B,C) Profile of (B) mouse- and (C) primate-specific maternal effect genes over developmental time in all species. (D–G) Gene ontology (GO) term enrichment analyses for zygotic transcripts (FPKM >10) in (D) primate, (E) human, (F) marmoset and (G) mouse. (H) Combined *Z*-scores of PRC1 and PRC2 components over seven developmental stages including 2-cell embryos.

**Fig. S3:**
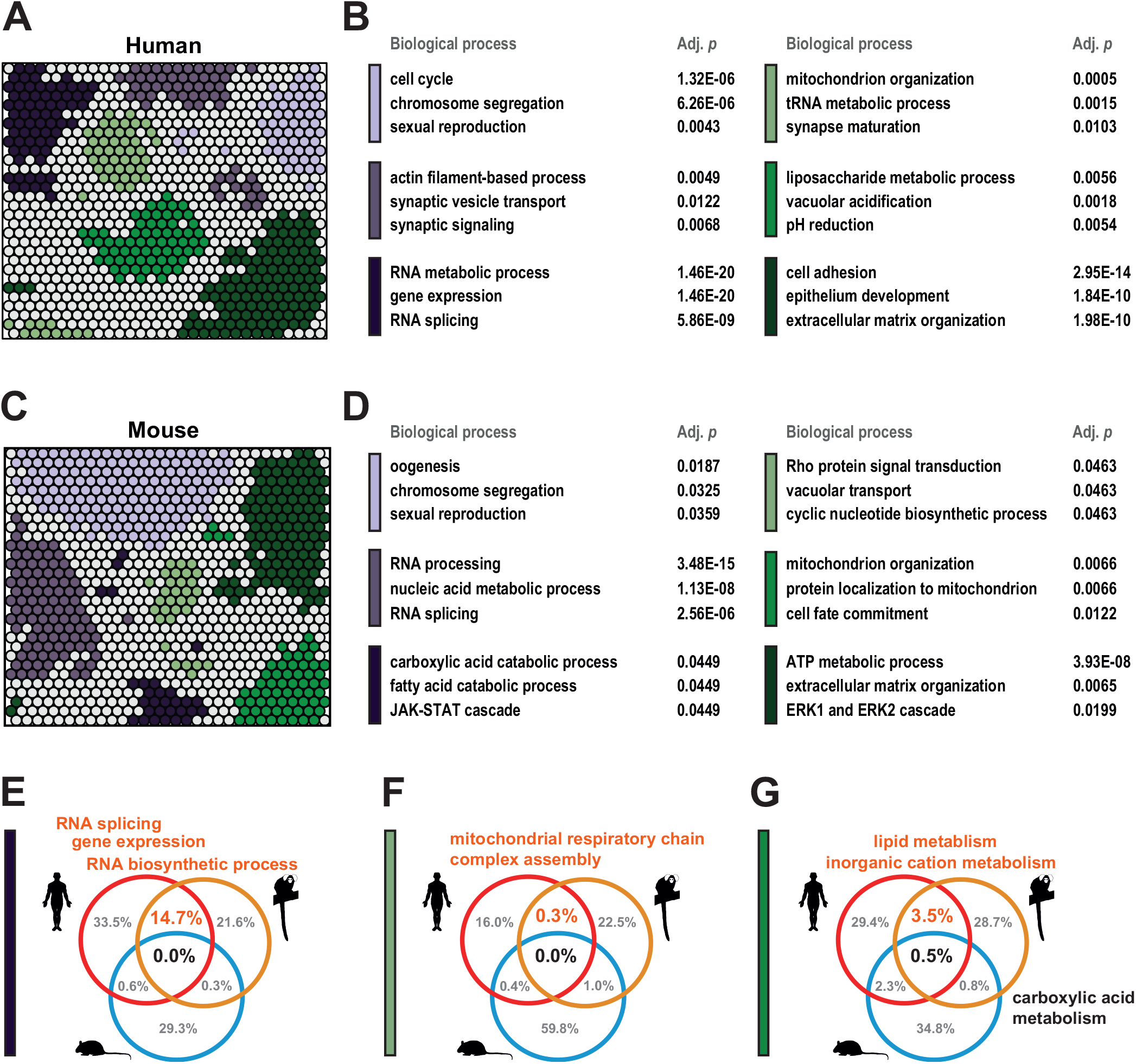
(A) Self-organizing map (SOM) of human embryo transcriptomes. Clusters (*Z*-score >1.5) are coloured by stage specificity. (B) Biological processes enriched for human SOM clusters. C) SOM of mouse embryo samples. (D) GO processes enriched in mouse SOM clusters. (E–G) Significantly enriched (*p* <0.05) biological processes in human, marmoset and mouse at the 8-cell (E), compacted morula (F) and early ICM (G) stages.

**Fig. S4:**
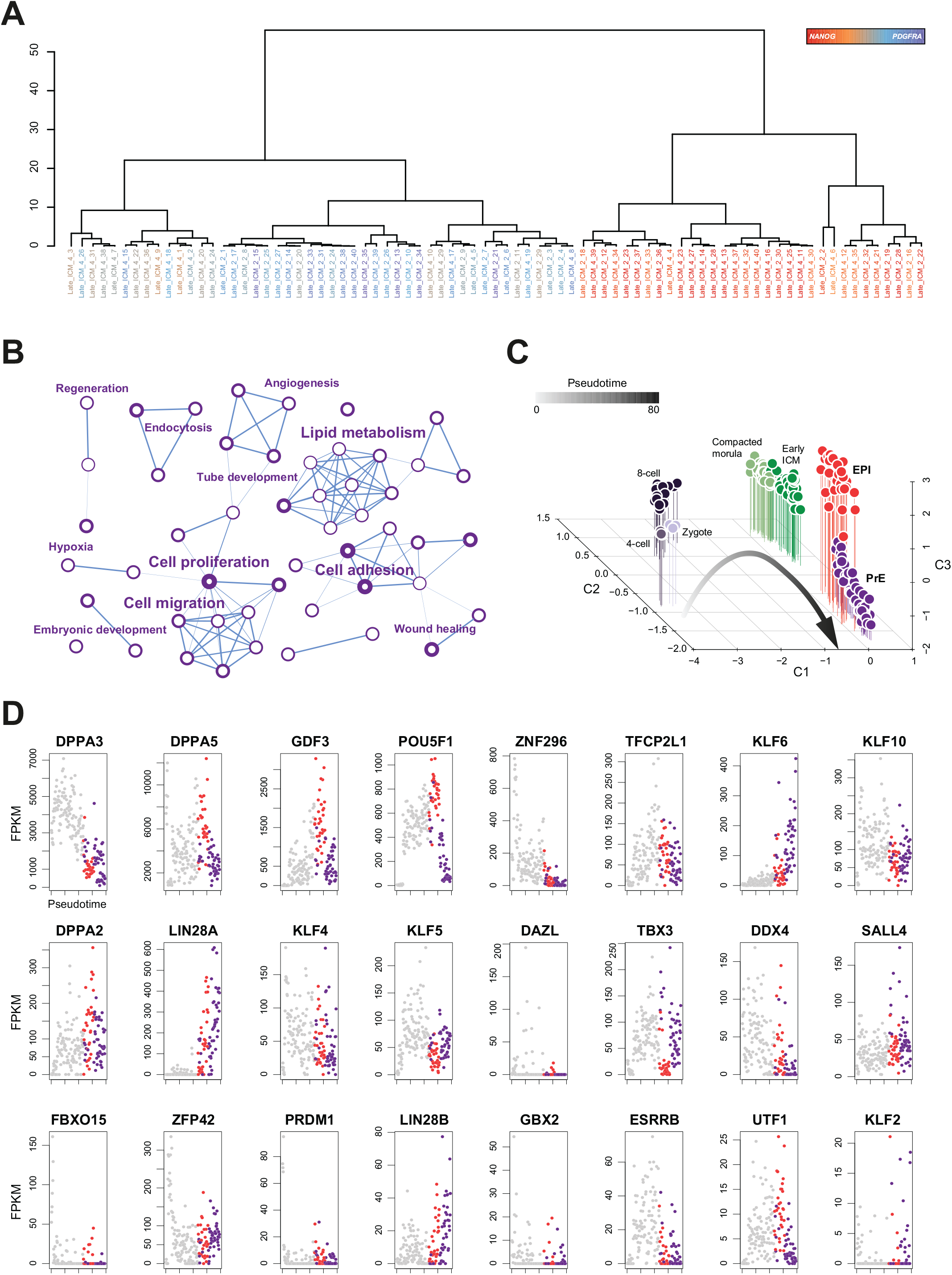
(A) Dendrogram of marmoset late ICM cells based on the first principal component of the PCA in Fig. 4B. (B) Enrichment map of the top 50 biological processes (*p* >0.05) based on absolute fold change >0.5 between PrE and EPI. (C) Individual component analysis (ICA) of embryo stages for derivation of developmental pseudotime. (D) Absolute expression of selected pluripotency and germ cell markers, ordered by pseudotime.

**Fig. S5:**
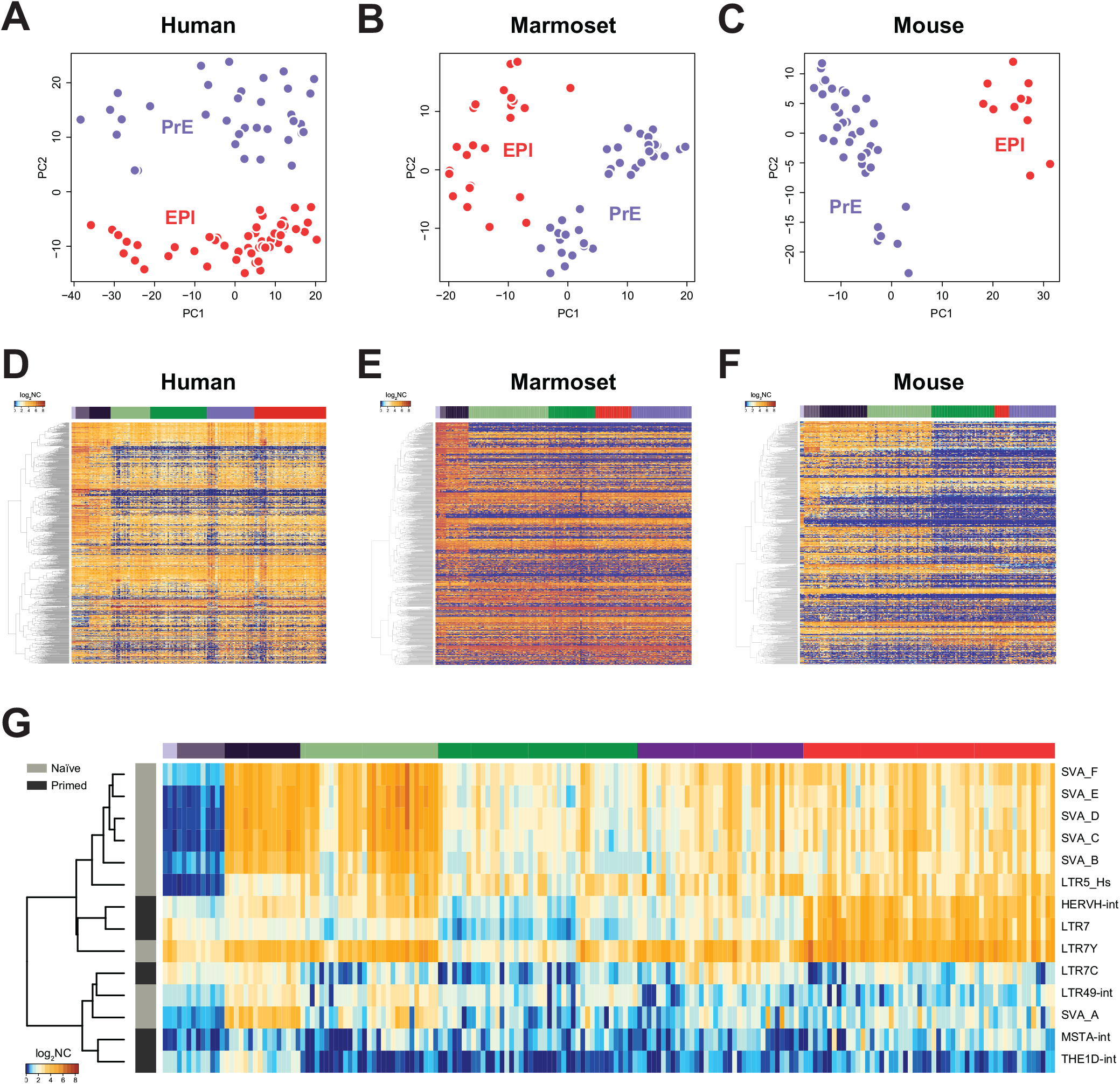
(A–C) PCA of human (A), marmoset (B) and mouse (C) EPI and PrE cells (log_2_ FPKM >0.5 and logCV^2^ >1). (D–F) One-way hierarchical clustering of averaged expression of transposable element families detected in human (D), marmoset (E) and mouse (F) embryos. (G) One-way hierarchical clustering of averaged expression of transposons associated with naïve and primed human PSC as defined in (Theunissen et al., 2016).

**Fig. S6:**
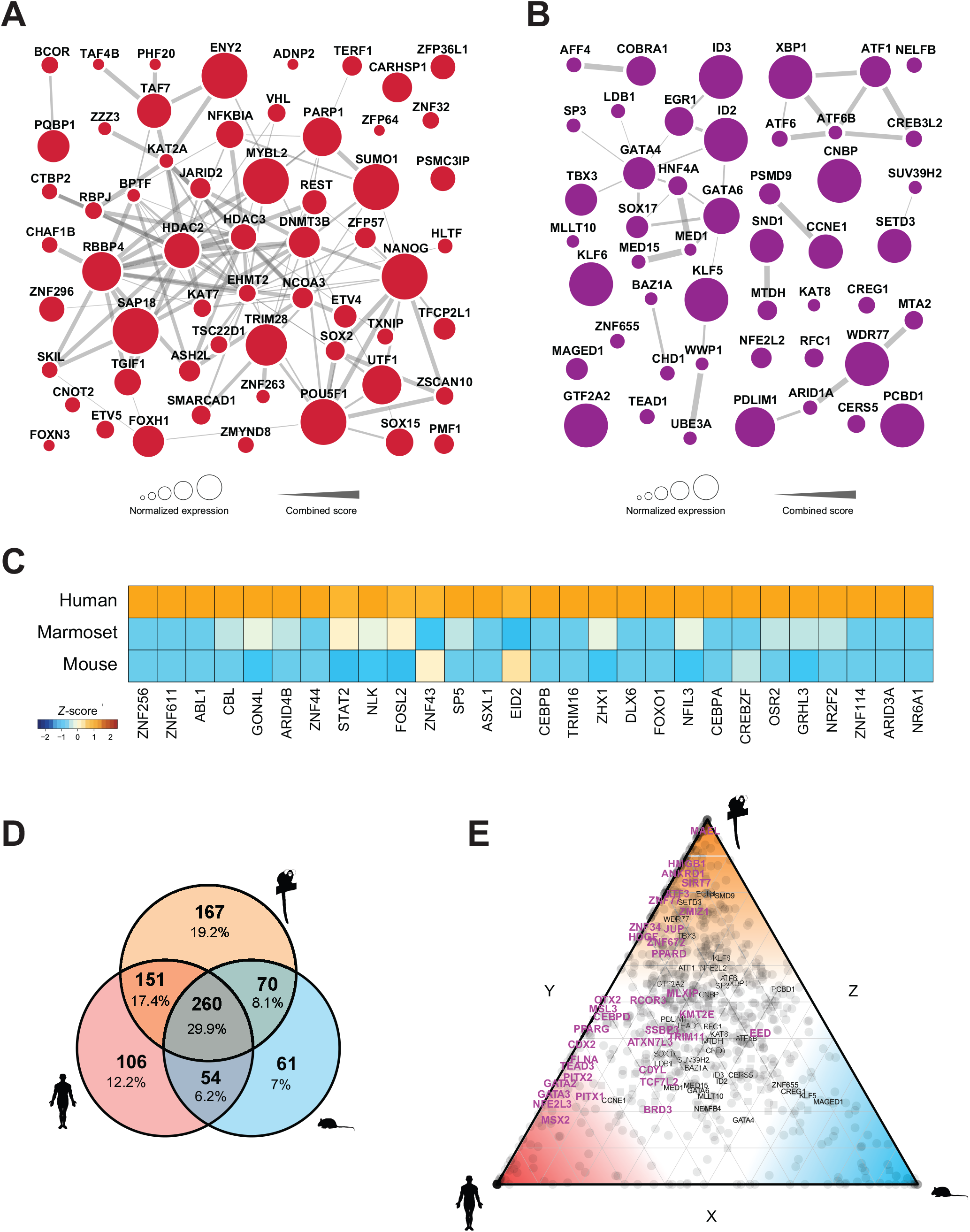
(A,B) Protein-protein interaction network of conserved EPI (A) and PrE (B) transcription factors in human, marmoset and mouse. Genes in (A) are derived from the analysis in Fig. 6A, and those in (B) from Fig. S6D above. Node sizes are scaled to normalised expression in human and marmoset; edges are derived from the STRING database. (C) Human transcription factors specific to primitive endoderm. (D) Intersection of transcription factors specific to human, marmoset and mouse PrE (FPKM >5 in PrE and not significantly (*p* >0.05) upregulated in EPI). (E) PrE-enriched transcription factors (circles) and chromatin remodelling factors (squares) between human, marmoset and mouse. Axes show the relative fraction of expression in the EPI between mouse and human (*x*), human and marmoset (*y*), and marmoset and mouse (*z*).

**Fig. S7:**
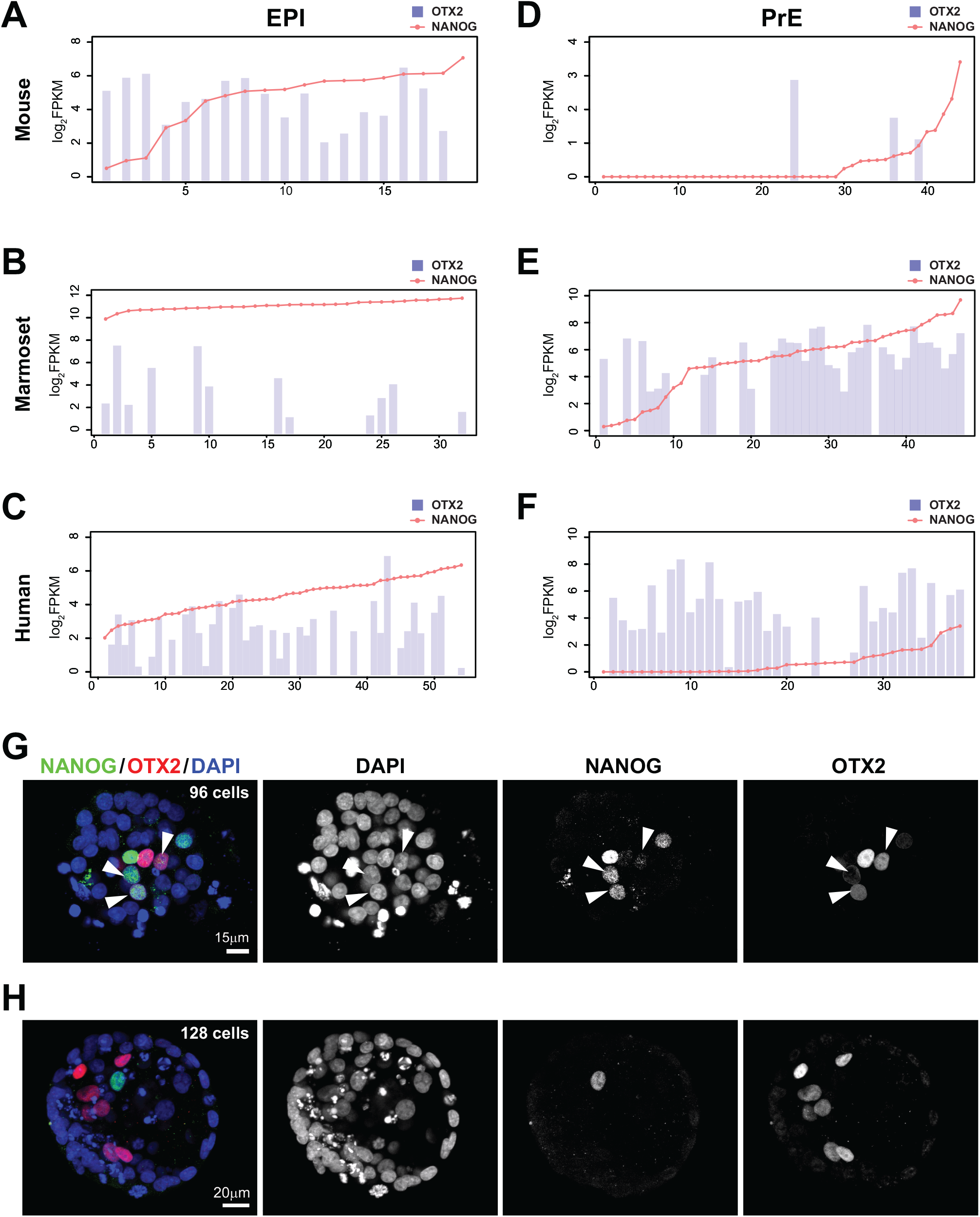
Plots generated by the web resource showing expression of *OTX2* in relation to *NANOG* in each species. (A–F) *OTX2* and *NANOG* mRNA levels in log_2_ FPKM for EPI in mouse (A), marmoset (B) and human (C), and PrE in mouse (D), marmoset (E) and human (F). (G,H) Confocal immunofluorescence sections of NANOG, OTX2 and DAPI in early (G) and late (H) human blastocysts. (G) Co-localisation of NANOG and OTX2 (arrows) is observed in cavitating embryos with intermediate protein abundance. (H) Mutually exclusive expression of NANOG and OTX2 is evident in later-stage embryos.

## Table Legends

**Table S1:** Libraries and coverage statistics for single-cell RNA-seq samples from common marmoset embryos, spanning zygote to late preimplantation blastocyst stages.

**Table S2:** Annotated orthologs for human, marmoset and mouse transcriptomes with average expression in preimplantation embryo stages.

**Table S3:** Differential expression of genes detected in EPI and PrE lineages for human, marmoset and mouse.

**Table S4:** Average gene expression in six preimplantation embryo stages for human, marmoset and mouse.

**Table S5:** Average expression of transposable elements annotated by family and class in six preimplantation embryo stages including EPI and PrE for human, marmoset and mouse.

**Table S6:** Average gene expression in six preimplantation embryo stages including EPI and PrE for human, marmoset and mouse, annotated for transcription factors, co-factors and chromatin modifiers.

**Table S7:** Classes and families of transposable elements for human, marmoset and mouse.

